# Small molecule mediated stabilization of PP2A modulates the Homologous Recombination pathway and potentiates DNA damage-induced cell death

**DOI:** 10.1101/2022.06.10.495657

**Authors:** Rita A. Avelar, Amy J. Armstrong, Gracie Carvette, Noah Puleo, Riya Gupta, Jose Colina, Peronne Joseph, Alex Sobeck, Caitlin M. O’Connor, Agharnan Gandhi, Michele L. Dziubinski, Daniel Shanhuai Ma, Steven Waggoner, Kristine Zanotti, Christa Nagel, Kimberly Resnick, Sareena Singh, Daffyd Thomas, Stephanie Skala, Junran Zhang, Goutham Narla, Analisa DiFeo

## Abstract

High-Grade Serous Carcinoma (HGSC) is the most common and lethal ovarian cancer subtype. PARP-inhibitors (PARPi) have become the mainstay of HGSC targeted therapy, given that these tumors are driven by a high degree of genomic instability and Homologous Recombination (HR) defects. Nonetheless, only ∼30% of patients initially respond to treatment, ultimately relapsing with resistant disease. Thus, despite recent advances in drug development and increased understanding of genetic alterations driving HGSC progression, mortality has not declined, highlighting the need for novel therapies. Using a Small Molecule Activator of Protein Phosphatase 2A (PP2A) (SMAP-061), we investigated the mechanism by which PP2A stabilization induces apoptosis in Patient-Derived HGSC cells and Xenograft (PDX) models alone or in combination with PARPi. We uncovered that PP2A genes essential for transformation (B56α,B56γ and PR72) and basal phosphatase activity (PP2A-A and -C) are heterozygously lost in the majority of HGSC. Moreover, loss of these PP2A genes correlates with worse overall patient survival. We show that SMAP-061 stabilization of PP2A inhibits the HR output by targeting RAD51, leading to chronic accumulation of DNA damage and ultimately apoptosis. Furthermore, combination of SMAP-061 and PARPi leads to enhanced apoptosis in both HR-proficient and -deficient cells and in patient-derived xenograft models. Our studies identify PP2A as novel regulator of HR and introduces PP2A activators as a potential treatment for HGSC tumors. Our studies further emphasize the potential of PP2A modulators to overcome PARPi insensitivity, given that targeting RAD51 has presented benefits in overcoming PARPi-resistance driven by BRCA1/2 mutation reversions.

## Introduction

High Grade Serous Cancer (HGSC) ranks as the 5^th^ leading cause of cancer related deaths in women and is the most common and lethal subtype of all female gynecological malignancies, accounting for ∼60% of all ovarian carcinomas(1,2). The American Cancer Society predicts that this year, 21,410 new cancer cases will be diagnosed and about 14,000 women will die of this disease. HGSC is most commonly diagnosed as late-staged metastatic cancer for which treatment options still rely on aggressive surgical resection as a single or a combinational approach with platinum-based chemotherapies(3). Nevertheless, the success rates remain highly discouraging (∼19% 5-year survival)(4) and acquired chemoresistance is still considered the main factor in tumor relapse and metastasis(5). Therefore, there is an urgent need for the development of novel and more efficient therapies for the treatment of HGSC. Recent advances in the targeted therapy field have introduced the use of PARP inhibitors (PARPi) for the treatment of ovarian and breast cancers, showing great promise clinically(6–8). PARPi comprise of small molecules that target the Poly [ADP-ribose] polymerase 1 (PARP1) to prevent it from repairing single strand DNA (ssDNA) breaks, resulting in the formation of double strand breaks (DSBs)(9). In the absence of functional PARP1, cells with HR deficiencies accumulate DNA damage leading to cell death in a process known as synthetic lethality. PARPi have been widely successful in treating HGSC patients that carry HR deficiencies, most commonly germline and/or somatic mutations in either BRCA1 or BRCA2 (BRCA1/2). However, when HGSC tumors are BRCA1/2 wildtype (or HR-proficient), PARPi dependent DSB formation can be resolved by the functional HR machinery, thus preventing cell death, and promoting cancer cell survival. Therefore, finding targeted therapies that can treat tumors that are no longer sensitive to PARPi or prevent acquired resistance to PARPi are of high priority.

Protein Phosphatase 2A is a large family of Serine/Threonine phosphatases consisting of three main subunits – PP2A-A, PP2A-B and PP2A-C – that form an active trimeric holoenzyme structure capable of dephosphorylating substrate proteins(10,11) (Fig. 1A). The scaffold (PP2A-A) subunit is a highly flexible protein that serves as the main platforming surface for all components to bind and closely interact(12). The catalytic (PP2A-C) subunit provides the phosphatase with its enzymatic capability. Finally, the regulatory (PP2A-B) subunit grants the phosphatase enzyme substrate specificity, and include 16 different subunits that are known to have unique target affinities(13). PP2A has been found to be impaired in the majority of human cancers, as its main role is tumor suppressive by acting as a master regulator of multiple essential cellular processes, counteracting oncogenic signals that drive malignancy(13–16). MAPK(11,17,18) and Wnt(18,19) pathways have been extensively studied as direct targets of PP2A dephosphorylation, showing that inhibiting these pathways presents promising antitumor activities. Most recently, the impact of PP2A on DNA Damage Response (DDR) and Homologous Recombination (HR) has also been established(20–24). Multiple PP2A heterotrimers were found to be essential in directing successful HR-dependent DSBs repair(24) and the BRCA2-PP2A complex formation/stability was established to be required for DNA repair efficiency by the HR pathway in human cells(25). Therefore, PP2A seems to represent an attractive therapeutic target for the treatment of human cancers that are driven by a high degree of Genomic Instability (GI), including HGSC. Given this information, developing new molecules that seek to directly stabilize, modulate, and selectively guide PP2A against oncogenic proteins that drive cancer development and progression holds potential clinical value.

**Figure 1.**
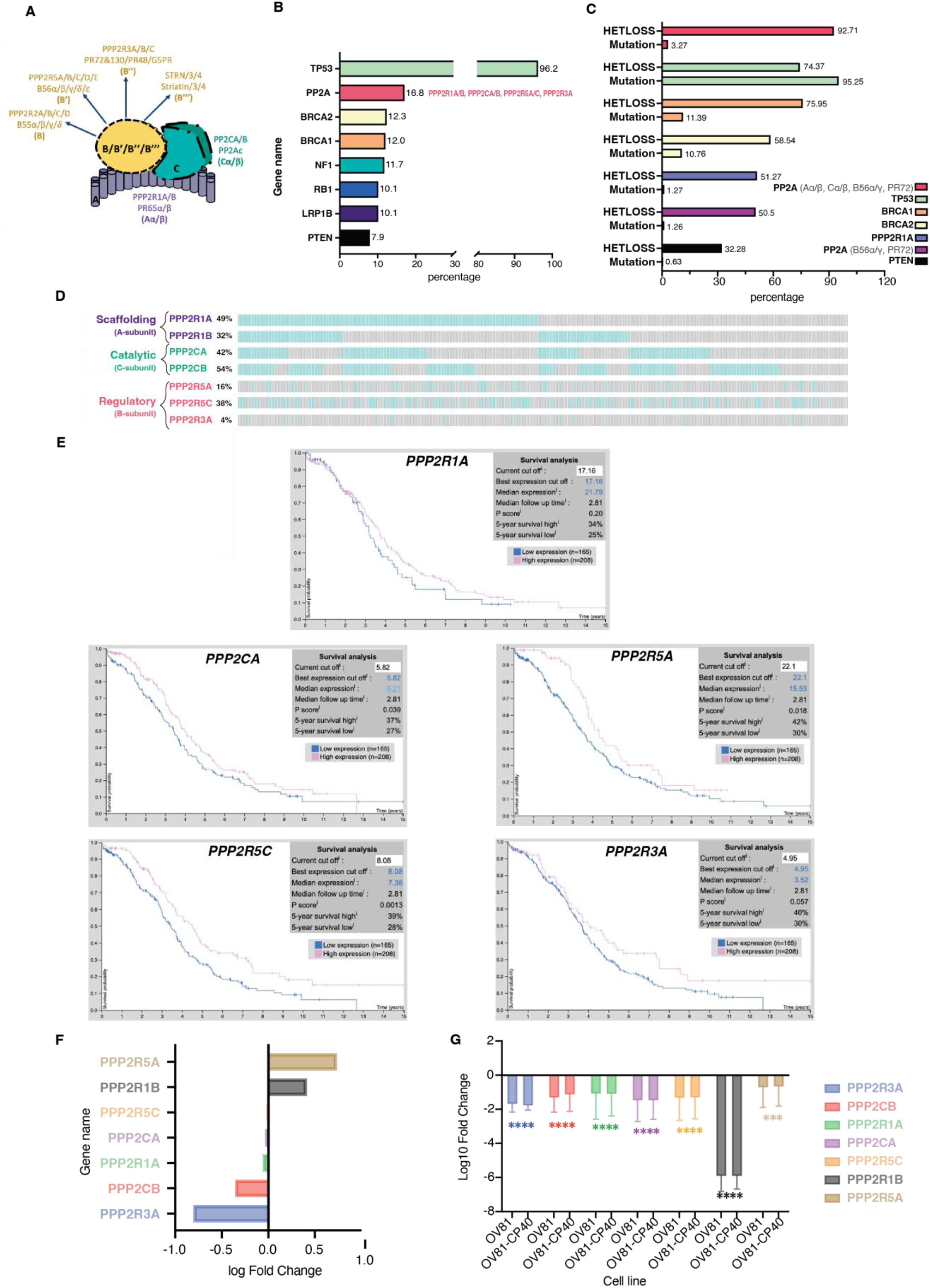
Dysregulation of PP2A is a common event in HGSC. A) Schematic of the PP2A heterotrimeric structure, representing all possible combinations of the scaffolding (A), regulatory (B) and catalytic (C) subunit proteins. B) Top 8 most common altered genes in HGSC and respective genetic alterations frequency (mutation, amplification and/or deletion). “PP2A” group includes PP2A-Aα/β, PP2A-Cα/β, PP2A-B56α/γ and PP2A-PR72). Data adapted from cBioPortal. C) HGSC genes alteration type specificity: Heterozygous/shallow deletion (Hetloss) or mutation; and respective frequencies (%). Data adapted from cBioPortal. D) Individual patient tumor data analysis showing frequency of heterozygous loss status for the PPP2R1A, PPP2R1B, PPP2CA, PPP2CB, PPP2R5A, PPP2R5C, PPP2R3A genes. Co-occurrence profiles are also shown. Each bar represents an individual tumor, for which grey signifies negative and blue positive for Hetloss. E) Kaplan-Meier plots summarizing results from correlation analysis between mRNA expression level and patient survival for PP2A genes: PPP2R1A, PPP2CA, PPP2R5A, PPP2R5C, PPP2R3A. Patients were divided according to expression levels into one of two groups “low” (under cut off on table) or “high” (over cut off on table). X-axis shows time for survival (years) and y-axis shows the probability of survival, where 1.0 corresponds to 100%. <u>Data source:</u> Protein Atlas. F) Log fold change of transcriptional expression of PP2A subunits between HGSC cell lines vs. FTSEC cell lines (FT246, FT33, FT194). <u>Source:</u> Elias *et al* 2016^44^. G) Log_10_ Fold Change of fragments per kilobase of exon per million mapped fragments (FMPK) of the previously assessed PP2A genes (Fig. 1D) comparing expression of the FT246 non-malignant with the OV81 and OV81-CP40 cell lines. Data presented as the mean ± SD (n=3), (unpaired Student T-tests, comparing OV81 and OV81-CP40 relative to FT246, ***p<0.005, ****p < 0.0001).

Targeting PP2A has been considered challenging given its promiscuity in target selection; therefore, finding a single molecule that activates an entire family of phosphatases has not been feasible. Recently, a small molecule SMAP-061 (or DT-061, as referenced by Leonard *et al*.) has been shown by several groups to act as a molecular glue which fits in a binding pocket created through the interaction between PP2A-A scaffolding subunit, the PP2A-C catalytic subunit and a PP2A-B subunit to effectively modulate PP2A holoenzymes(26), driving downstream dephosphorylation of essential oncogenic drivers such as c-Myc, AR, ERK, AKT. Cryo-EM studies suggested that the SMAP compounds preferentially recruit PP2A-B56α subunit to form a trimeric holoenzyme(26), however alternative mechanisms have also been proposed wherein SMAP-061 specially stabilizes the B55α-PP2A heterotrimer in blood cancers. Despite the slight contradiction among studies, multiple groups consistently agree that studying these SMAP molecules is essential to further evaluate the therapeutic potential of PP2A modulation, and further explore the mechanisms by which they exert their potent anticancer properties(26–30). SMAP-061 has potent anticancer properties both *in vitro* and *in vivo* resulting in reduced tumor burden in various human cancer models without any issues with tolerability, even in long-term dosing studies *in vivo*(26–29,31,32). Furthermore, SMAP-061 and other established molecules mediators of PP2A(30) have been shown to hold immense potential for the treatment of a wide range of human cancers in pre-clinical studies using xenograft, genetically engineered and orthotopic mouse models, including prostate(27), breast(31), lung(28,29), glioblastoma(32), etc., with efficacy being linked to the dephosphorylation of oncogenic substrates.

Here, our research studies began by defining the expression of PP2A in HGSC, we found that greater than 90% of HGSC tumors display heterozygous loss of at least one of the PP2A genes essential for malignant transformation and/or holoenzyme formation, second only to p53. In addition, we uncover that HGSC tumors with decreased expression of these genes correlate with worse overall survival rates. We report that this recurrent genetic perturbation in PP2A can be leveraged therapeutically using SMAP-061 in HGSC HR-proficient or deficient. We show that treatment of HSGC cells with SMAP-061 can effectively reduce tumor burden by decreasing the expression and activity of multiple DDR and HR proteins. Furthermore, we confirmed that SMAP-061 induced RAD51 protein expression loss necessary for apoptosis and postulated that inhibiting RAD51 filament formation impairs the high-fidelity HR repair pathway and sensitizes HGSC cells to PARPi, ultimately leading to induced synthetic lethality in this malignant context. Lastly, we find that SMAP-061 works additively or synergistically with Olaparib in Patient-Derived (PD) HGSC models with varying sensitivities to PARPi. In sum, our data suggests that SMAP-061 represents a unique opportunity to expand the patient population that can benefit from PARPi, no longer having to rely only on tumors with a HR deficiency (HRD) profile.

## Results

### Loss of PP2A is a frequent event in HGSC

Due to the heterogenous nature of HGSC, it has been challenging to validate and functionally identify putative genes that drive its tumorigenesis, even though HGSC has been widely defined by a high degree of GI. Approximately 50% of all HGSC are known to have genetic alterations (mutation, amplification and/or deletion) in the HR pathway, with *TP53, BRCA1/2* and *RB1* listed at the top (Fig. 1B). The PP2A family of phosphatases has been found to be essential to tightly regulate the DDR and HR pathways, directly dictating phosphorylation events of multiple downstream targets(22,33). Furthermore, Karst *et al* has demonstrated that loss of PP2A is essential for the transformation of the Fallopian Tube Secretory Epithelial Cells (FTSEC or FT) model(34), further suggesting an important role for PP2A in HGSC carcinogenesis. Interestingly, opposite to other female gynecological cancers such as serous endometrial cancers(35–37), literature shows that PPP2R1A mutational burden for ovarian cancers are rare across most histological subtypes(38), suggesting that the inactivation mechanism of PP2A in this tissue type must be accomplished in a different manner. Nevertheless, gene alterations to PP2A and their direct link and impact on HGSC development and progression have not yet been fully explored. Thus, we first set out to define the differential expression of PP2A genes that have been shown to drive tumorigenesis and correlate it with clinical outcomes in HGSC. We identified multiple different genes in the PP2A family that were altered to a higher extent than the majority of the well-established and extensively studied HR genes in HGSC (Fig. 1B and Fig. 1C). According to the TCGA, PPP2R1A is found to be heterozygously deleted/lost (HetLoss) in about 50% of all HGSOC, and at least 1 out of the 3 B-subunits previously established to be essential for malignant transformation (B56α, B56γ and PR72)(39–41) is also found to be Hetloss in ∼50% of all HGSOC (Fig. 1C). When we looked at the probability of the minimal essential genes for PP2A enzymatic/phosphatase basal activity (ie, PP2A-A and PP2A-C, as the dimer alone still renders baseline activity, despite lack of specificity) and the B subunit genes known to have transformative potential (B56α, B56γ and PR72), we found that these genetic alterations comprise for the majority of Hetloss percentage found in all HGSC tumors (Fig. 1C PPP2R1A/B, PPP2CA/B, PPP2R5A/C, PPP2R3A) - 92.7%. However, these Hetloss alterations are not mutually exclusive and certain patients may harbor these shallow deletions in more than one of these PP2A genes at a time (Fig. 1D) (for instance, the first patient harbors Hetloss in PPP2R1A/B, PPP2CA/B, and PPP2R5C genes). All other B-subunits alterations percentages have also been individually analyzed (Table S1A). Interestingly, somatic mutations seem to be overall extremely rare (3.48%) (Fig. 1C and Table S1A), suggesting that optimizing PP2A heterotrimer formation and biasing the existing pools towards a more tumor suppressive holoenzyme structure seems to be a promising strategy to produce the optimal and desired anticancer effects in HGSC tumors. Given that further analysis shows that copy-number alterations proportionally correlate with their respective mRNA expression levels (Fig. S1A-S1F). We then proceeded in evaluating whether the mRNA expression for these specific genes correlate with patient outcome. We found low mRNA expression of PPP2R1A, PPP2CA, PPP2R5A, PPP2R5C and PPP2R3A correlates with decrease in patient survival rates (Fig. 1E). This data suggests that the expression levels of these specific PP2A genes directly impact patient survival. Individual patient tumor data also demonstrates that Hetloss is not mutually exclusive within the PP2A genes (Fig. S1G and Table S1B), and its co-occurrence is prevalent with similar combination patterns between the scaffolding, catalytic and regulatory subunits (Fig. S1H). Additionally, it has been previously established that PP2A-A is a haploinsufficient tumor suppressor gene both in cellular and *in vivo* model systems(39,42,43) further lending support to our idea that these highly recurrent alterations in HGSC might play pathogenic roles.

Together, these observations led to the hypothesis that the loss of specific PP2A genes results in attenuation of its tumor suppressive function, possibly promoting the development and progression of HGSC. Moreover, the fact that these genes seem to have an extremely low mutational burden, would indicate that selective deletion of specific pools of PP2A activity is critical for HGSC tumorigenesis. Thus, therapeutic reactivation and modulation of the remaining pool of PP2A heterotrimers might be sufficient to trigger HGSC cell death in cellular and *in vivo* model systems of this disease.

### SMAP-061 treatment induces apoptotic cell death in HGSC

Given that a large majority of HGSC tumors displayed heterozygous loss of at least one of the PP2A genes and loss of PP2A has been shown to be required for HGSC transformation^51^, we next wanted to investigate whether PP2A modulation using SMAP-061 would affect the viability of HGSC PD (Fig 2A, left and Table S2A) and isogenic cell lines (Fig. 2B, left and Table S 2B). We first confirmed that the subunits commonly lost in HGSC tumors were significantly reduced in the HGSC models compared with FT lines (Fig. 1F and Fig. 1G).(44) Interestingly, we found that all lines were sensitive to SMAP-061, with EC_50_ values ranging between 10 and 20µM, despite the wide range of responses observed with the treatment of cisplatin, which is the first-line chemotherapy for HGSC (Fig. 2A, right and Table S3A). To further assess whether cisplatin sensitivity and/or HR defects modified SMAP-061 response, we utilized isogenic cisplatin sensitive and resistant lines (Fig. 2B and Table S2B). OV81 and its Cisplatin Resistant (CR) pair OV81-CP40 showed very distinct responses to cisplatin given that these cells were grown in low cisplatin concentrations for 40 passages, however their response to SMAP-061 remained the same (Fig. 2B, left). The PEO-1 and its isogenic pair PEO-C4.2 showed similar responses to SMAP-061 however did not show resistance to cisplatin treatment most likely due to only being exposed to cisplatin for 2 passages. However, what is unique about this pair is that the parental PEO-1 cell line is HR-deficient given that it has a BRCA2 mutation 5193C>G(Y1655X) and PEO-C4.2 has acquired an BRCA2 reversion mutation thus making it HR-proficient^56^ (Table S2B), (Fig. 2B, right). Therefore, these findings show that SMAP-061 has similar potency despite differences in platinum sensitivities and HR status (Fig. 2B), as previously observed in the PD HGSC panel (Fig. 2A). To further investigate how the different HR efficiency profiles would impact and dictate the molecular response of HGSC lines to SMAP-061 treatment, we selected a PD HGSC line (OV81) and the isogenic HR-deficient and proficient cells (PEO-1 and PEO-C4.2) for our future experiments. To assess the long-term effects of SMAP-061 treatment on these cells we performed colony forming assays and found that SMAP-061 treatment resulted in significantly fewer colonies (Fig. 2C and Fig. 2D). Moreover, SMAP-061 treatment led to a significant induction of apoptosis in a time dependent manner, as indicated by the increase of cleaved PARP and cleaved Caspase 3 in the three HGSC cell lines (OV81, PEO-1 and PEO-C4.2) (Fig. 2E). Given that HGSC are derived from the fallopian tube epithelium, we wanted to further investigate whether these apoptotic effects would also be observed in the non-malignant precursor cells or if they were specific to their malignant counterparts. We found that the benign Fallopian Tube epithelial cell line 246 (FT246) was less sensitive to SMAP-061 when compared to HGSC cells (FT246 EC_50_: 22.97µM vs. OV81 EC_50_: 11.38µM, Fig. S3D) and apoptotic cues were not triggered when the same dose of SMAP-061 was used as in the cancer cells (Fig. 2E).

**Figure 2.**
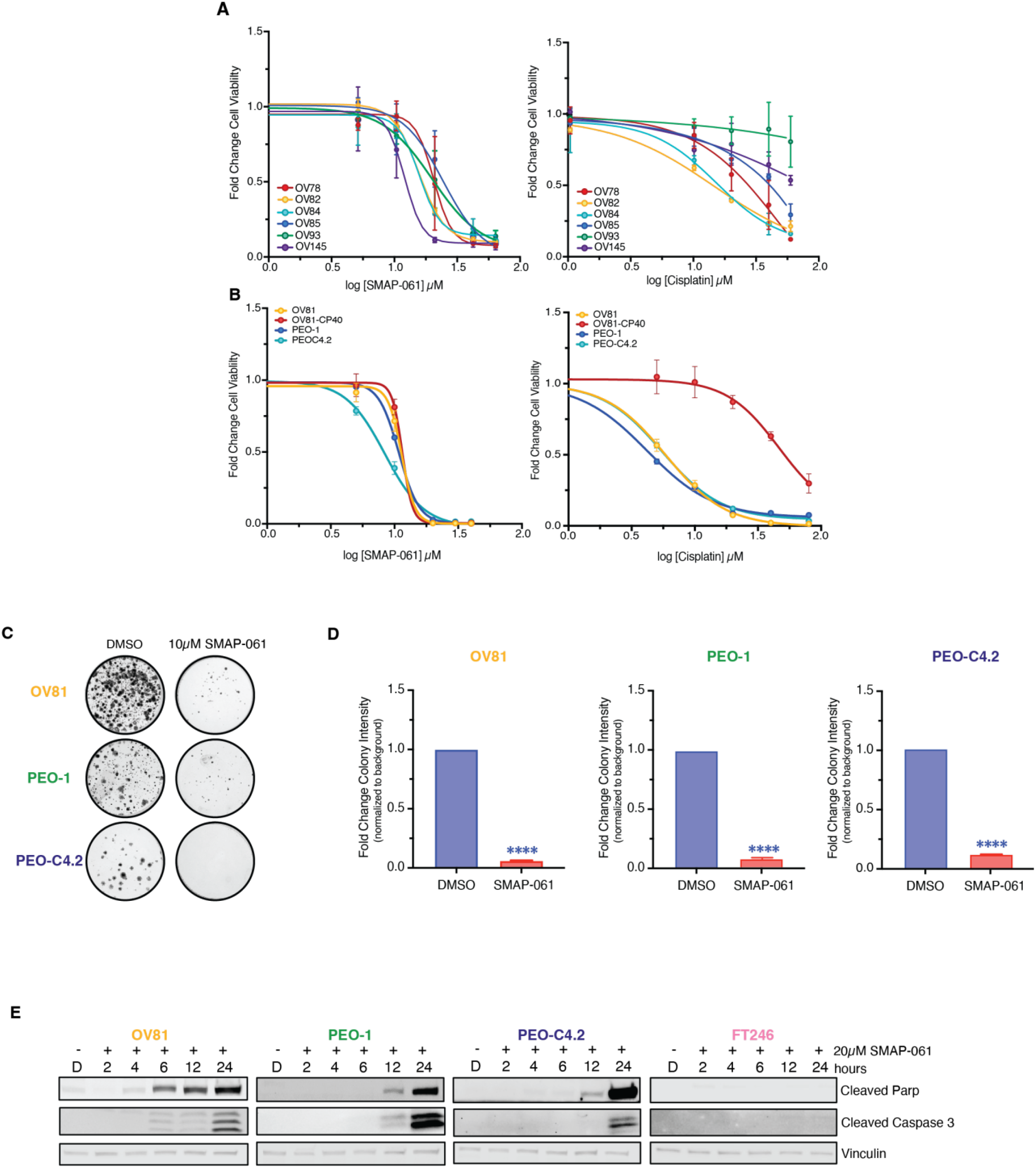
HGSC cells are sensitive to SMAP-061 and have similar responses independent of genetic background. A) A wide panel of HGSC PDX cell lines were exposed to increasing doses of either SMAP-061 (left) or cisplatin (right) treatment and cell viability was measured at 24 hours by MTT analysis, to generate an EC50 curve. Data is presented as the mean ± SD (n=3). B) OV81 and its cisplatin resistant pair, OV81-CP40, and two isogenic lines PEO-1 and PEO-C4.2, were treated with increasing doses of either SMAP-061 (right) or cisplatin (right) and cell viability was measured at 24 hours by MTT analysis, to generate an EC50 curve. Data presented as the mean ± SD (n=3). C) Clonogenic assay of OV81, PEO-1 and PEO-C4.2 cells treated with DMSO or 10µM of SMAP-061 for two weeks. D) Quantification of 2C. Data presented as the mean ± SD (n=3), (unpaired Student T-tests, comparing SMAP-061 treatment relative to DMSO, ****p < 0.0001). E) OV81, PEO-1, PEO-C4.2, and FT246 treated with DMSO or SMAP-061, and harvested at 2, 4, 6, 12 and 24 hours for western blot analysis of cell death markers (cleaved Parp and cleaved Caspase 3). Vinculin housekeeping gene used as loading control.

Together, this data suggests that stabilization and, ultimately, activation of PP2A leads to robust apoptotic effects in clinically and genetically diverse PD primary and established HGSC cells.

### SMAP-061-induced apoptosis is due to inhibition of HR and DDR signaling output and unresolved DNA damage

A hallmark of HGSC cells is their ability to withstand high levels of DNA damage, due to defects in HR and a high degree of genomic instability(45). These HR-deficient HGSC cells continue to rely on and benefit from the minimally functional HR pathway, in order to maintain sufficient levels of DNA repair that can sustain cell survival. Given that SMAP-061 can induce cell death in a wide-array of HGSC cell lines which are both HR-proficient and deficient and PP2A has been shown to potentially regulate several components of the HR pathway, we next wanted to examine whether this compound was affecting this signaling pathway (Fig. 3A). To do so, we analyzed the protein expression of multiple HR pathway family members after SMAP-061 treatment by western blotting as well as γH2ax foci by immunofluorescence (Fig. 3B-3E and Fig. S2A-S2L). SMAP-061 treatment increased γH2ax foci formation by 70% compared to DMSO control, after 12 hours of treatment (Fig. 3E), suggesting SMAP-061 promotes unresolved DNA damage accumulation. Interestingly, unlike what is commonly seen with DDR signaling in that the sensors proteins are induced first and then the transducer and effectors are activated to repair the DNA damage (Fig. 3A), we found that loss of RAD51 and WEE1 occurred significantly earlier (∼2 hours) and prior to the accumulation of γH2Ax expression, foci formation as well as p-RPA which was seen at ∼6-12 hours post-treatment (Fig. 3B-3D). As expected, the downstream phosphorylation and activation of direct targets such as ATM and CHK1/2 proteins also increased, as an attempt to fully activate HR and proceed to repairing the DNA insults. By remaining unresolved, such damage accumulation ultimately triggered apoptosis, as previously observed (Fig. 2E). Furthermore, there is a significant inverse correlation between RAD51 and γH2Ax expression for all three HGSC cell lines (Fig. 3F), indicating that the loss of RAD51 may be contributing to the accumulation of γH2ax. Interestingly, the observed phenotype is independent of HGSC BRCA genes’ status and HR genetic background, as all three cell lines show similar molecular signatures after treatment with SMAP-061, except for the RAD51 degradation and γH2ax accumulation, occurring in a more delayed pattern in PEO-C4.2 cells, which harbor a BRCA2 reversion mutation (Fig. 3B-3D). These results suggest that SMAP-061 could be inducing synthetic lethality through the regulation of crucial HR pathway proteins, thus leading to the accumulation of unresolved DNA damage and concomitant cell death (Fig. 3A).

**Figure 3.**
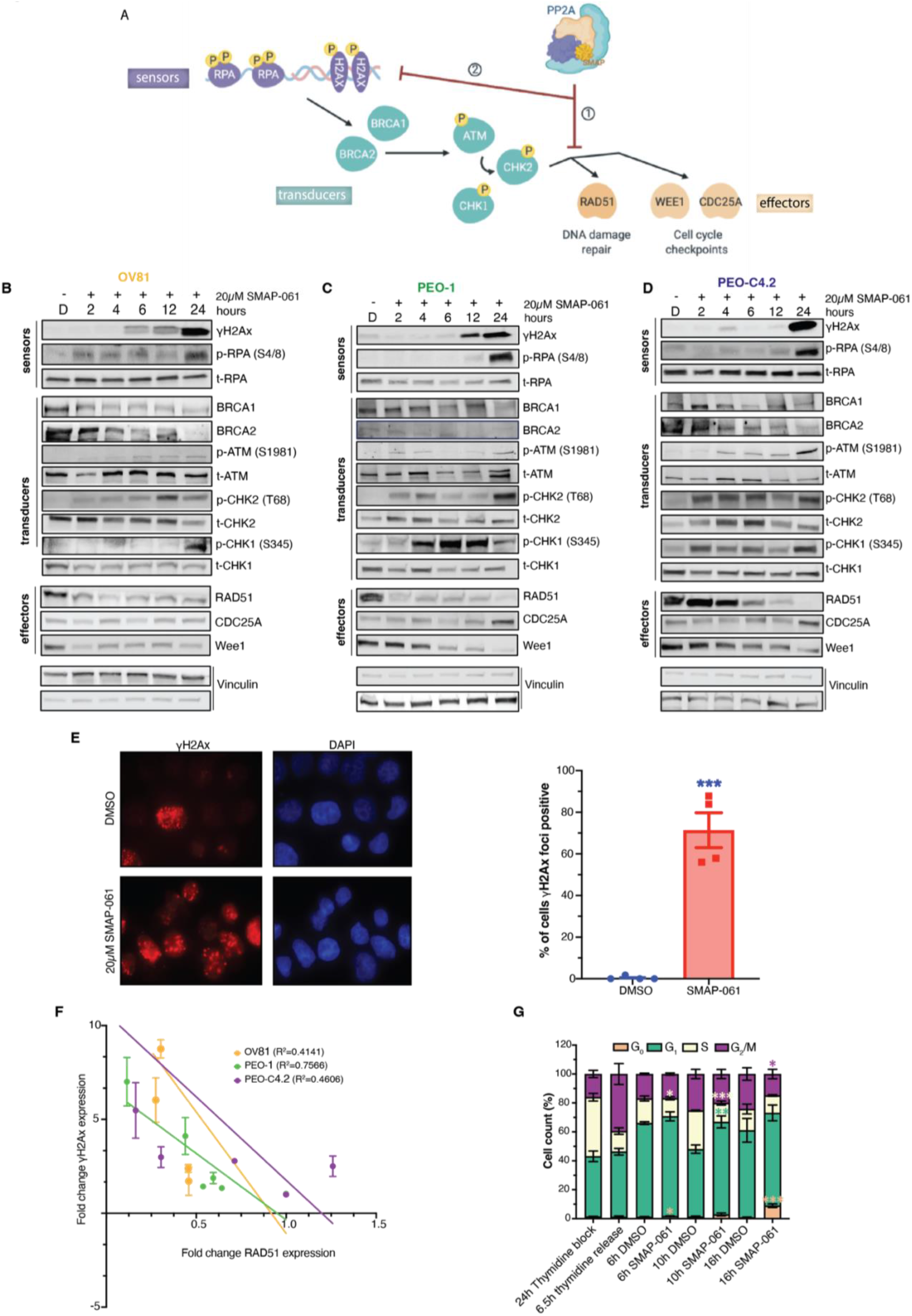
SMAP-061-induced baseline DNA damage accumulation is a result of the inhibition of HR signaling output, including DNA damage repair and cell cycle regulation mechanisms. A) Schematic of the HR pathway, representing the DNA **sensors** (RPA and H2Ax) that detect damage, the **transducers** (BRCA1/2, ATM and CHK1/2) that can amplify and transmit those damage signals to the **effectors** (RAD51, WEE1 and CDC25A), which ultimately regulate DNA damage repair and cell cycle checkpoint activity. Ultimately, when the HR output is successfully transmitted to the sensors by RAD51, HGSC baseline DNA damage is kept low and homeostasis is restored, allowing cells to survive long-term with inherent DNA errors due to Genomic Instability (GI). However, when SMAPs are added, PP2A gets activated, allowing for ① RAD51 expression to be inhibited and leading ② to the HR effectors’ function to be chronically impaired. This results in the incapacity of the cells to restore cell cycle progression and DNA damage repair, eventually dying to self-triggered apoptosis. Western blot analysis of B) OV81, C) PEO-1 and D) PEO-C4.2 cells after 2, 4, 6, 12 and 24 hours of SMAP-061 treatment evaluates the expression of **sensor, transducer** and **effector** proteins illustrated in the schematic of 3A. E) γH2Ax foci imaging comparing DMSO and 20µM SMAP-061 treated OV81 cells after 12 hours of incubation and their respective quantification. Data presented as the mean ± SEM (n=3), (unpaired Student T-tests, comparing SMAP-061 treatment group relative to DMSO, ***p < 0.001). F) Correlation analysis graph and R^2^ values comparing the expression of RAD51 with γH2Ax in OV81, PEO-1 and PEO-C4.2 during 2, 4, 6, 12 and 24 hours of SMAP-061 treatment. Data presented as the mean ± SEM (n=3). G) Statistical and quantification analysis of cell cycle flow (from Fig. S2O) for cells after incubation with 2mM thymidine for 24 hours (first bar), 6.5h thymidine released (to allow cells to reengage cell cycle progression) (second bar), and consecutive treatments with either DMSO or SMAP-061 for 6 hours (third and fourth bars), 10 hours (fifth and sixth bars) and 16 hours (seventh and eighth bars), respectively, using FlowJo. Data presented as the mean ± SD (n=3), (unpaired Student T-tests, comparing treatment groups to each other, for each stage of the cell cycle, *p < 0.05, **p < 0.01, ***p < 0.001).

### SMAP-061 treatment results in G_1_ cell cycle arrest and reduction of Cyclin D1 protein expression

Given that previous studies in multiple myeloma cells showed that RAD51 inhibition leads to similar increases in unrepaired DNA Damage due to a G_1_ cell cycle arrest(46), we next wanted to assess whether SMAP-061 could also directly affect cell cycle progression. After synchronizing HGSC cells using thymidine to allow them to undergo a full cycle, our data revealed that SMAP-061 treatment induced a similar G_1_ cell cycle arrest, followed by a progressive accumulation of an apoptotic population (G_0_) (Fig. 3G and Fig. S2M-S2N). Specifically, cells treated with DMSO after 10 hours post thymidine release, were able to normally transition into S-phase (**S-phase**: ∼25%) (Fig. S2N) however, HGSC cells treated with SMAP-061 were unable to transition into S-phase after 10 and 16 hours of treatment (**S-phase**: ∼11% and 12%, respectively) (Fig. 3G and Fig. S2N, top). These results suggest that SMAP-061 treated cells were continuously halted at G_1_ cell cycle checkpoint and unable to reengage back or progress through the different stages of cell cycle (Fig. S2N, bottom). Furthermore, SMAP-061 treated cells showed that the inability to leave G_1_ eventually lead to their accumulation in sub G_1_, indicating that these cells were undergoing cell death. We then analyzed the expression of Cyclin D1, an essential cyclin that functions as G_1_ cell cycle checkpoints gatekeeper. Western blots showed that Cyclin D1 expression decreases as early as 2 hours post SMAP-061 treatment (Fig. S2O-S2Q), possibly explaining the previously observed arrest phenotype. The expression levels of different cyclins (Cyclin B1 and E) or other cell cycle proteins were not affected to the same degree as D type cyclins in SMAP-061 treated cells, indicating that SMAP-061 may be specifically targeting E2F responsive genes (Cyclin D1 RAD51, BRCA1, & BRCA2) which are known to decrease in G_1_ arrested cells. Moreover, mitotic specific proteins could not reinstate protein turnover over time, given the downregulation of PLK1, Cyclin B1 and Securin (Fig. S2O-S2Q).

In sum, these results indicate that SMAP-061 treatment inhibits cell cycle transition by specifically impairing the capacity of HGSC cells to progress through G_1_. More importantly, by reducing Cyclin D1 expression, SMAP-061 prevents cells from transitioning into S-phase, where most HR proteins are commonly expressed. This process slowly leads to a chronic accumulation of γH2Ax and ultimately induces programmed cell death, due to synthetic lethality.

### SMAP-061 treatment potentiates PARPi effects *in vitro* and leverages synthetic lethality

Given that impact of SMAP-061 on regulating the expression of HR proteins, we next sought to assess whether combining PP2A activators with PARP inhibitors could effectively induce synthetic lethality in patients that lack HRD and/or potentiate the effects of PARPi in patients that developed PARPi resistance. PARP inhibitors are a first-line maintenance treatment option for HGSC patients, after treatment with platinum-based therapies. However, the mechanism behind response and sensitivity of tumors to PARPi before and after they become CR, has not been fully explored. Interestingly, the cisplatin-naïve OV81 and PEO-1 cell lines displayed a greater sensitivity to the PARPi Olaparib when compared to their respective CR pair (Fig. 4A-4D, Table S3B), potentially indicating an overlap in resistance mechanisms between PARP inhibitors and cisplatin. To evaluate whether SMAP-061 could potentiate PARPi response in the resistant models, we performed cell viability assays using increasing doses of SMAP-061 and Olaparib, as single agents or in combination, in both platinum naïve and resistant cell lines. Both the OV81 (Fig. 4A and Fig. 4C) and PEO (Fig. 4B and Fig. 4D) pairs showed increased cell death when the two drugs were combined comparatively to each agent alone, decreasing SMAP-061 EC_50_ more than two-fold (Table S3B). This was further confirmed by conducting an isobologram analysis and determining the Combination Index (CI) for SMAP-061 and Olaparib (Fig. 4E) in our HGSC lines. We found that the CI for all cell lines was <1, which indicates that the combination of the two drugs induces a synergistic effect. To determine whether the synergistic impact of SMAP-061 was specific to only PARP inhibitors, we then examined SMAP-061’s impact on how the cells respond to platinum-based therapies, which remain as the first-line treatment option for HGSC (Fig. S3A-S3C and Table S3C). Interestingly, despite SMAP-061 not sensitizing CR cells to cisplatin treatment, it was able to induce synergistic cell death in the cisplatin-naïve PEO-1 cells (CI=0.6) (Fig. S3B and Table S3C). In addition, SMAP-061 sensitized non-malignant FT246 cells to PARPi, however the doses required to induce cell death were significantly higher than in cancer cells (Fig. 2E, Fig. S3E and Fig. S3F). We also performed colony forming assay to evaluate long-term proliferation and survival effects of the combination drug treatment. We observed that in PEO-1 and PEO-C4.2 the combination treatment of SMAP-016 and PARPi was significantly more effective than either drug alone, however no synergy or additivity was observed in the OV81 cell line at the selected doses (Fig. 4F and Fig. S4A). Furthermore, protein analysis of cellular death (cleaved Parp and cleaved Caspase 3) and DNA damage markers (γH2Ax) by western blot further reflects these additive and synergistic outcomes in a dose (Fig. 4G, Fig. S4B, Fig. S4D, and Fig. S4F) and a time (Fig. 4H, Fig. S4C, Fig. S4E, and Fig. S4G) dependent manner.

**Figure 4.**
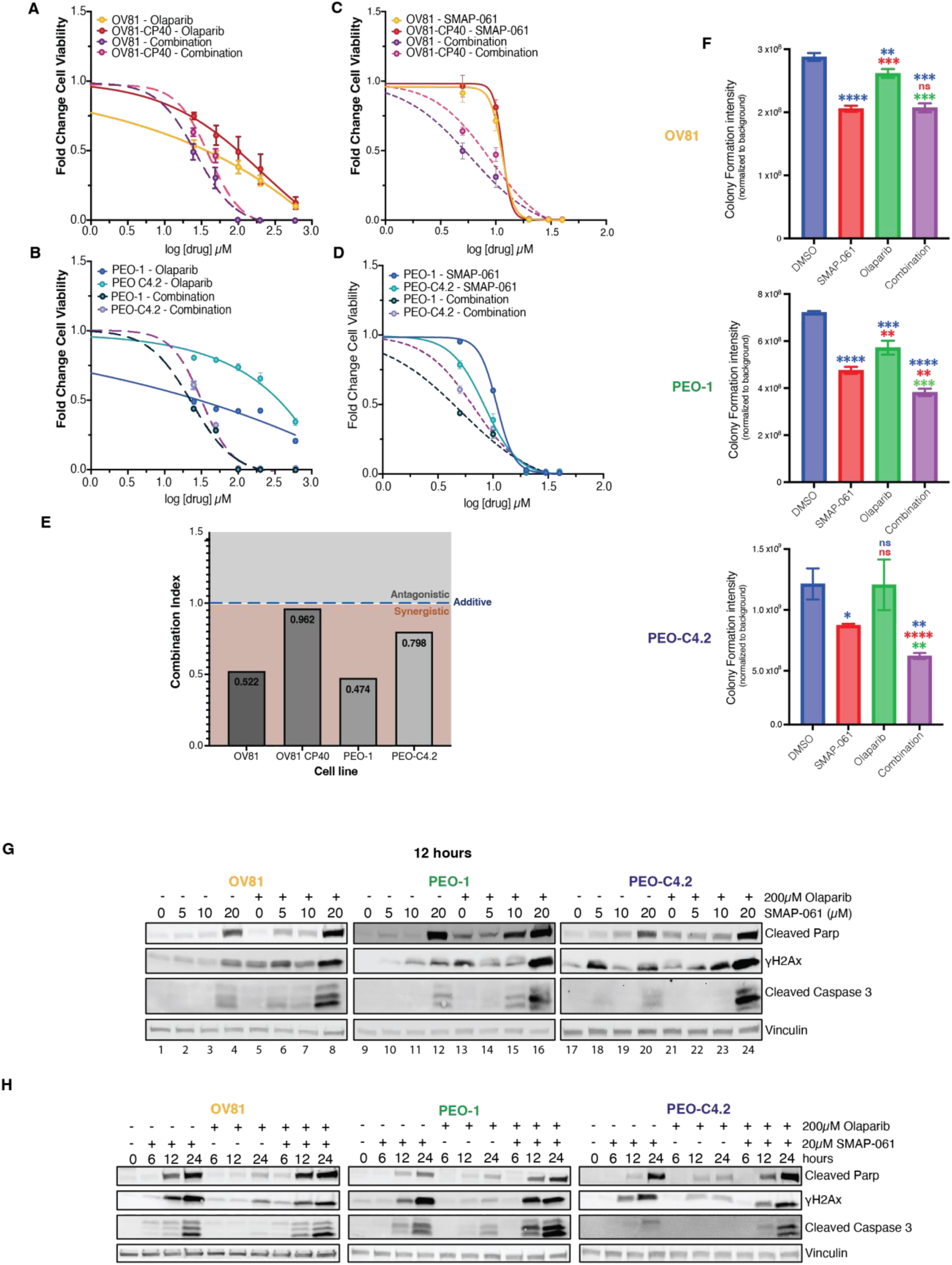
SMAP-061 shows synergistic efficacy in targeting HGSC for cell death when used in combination with PARPi *in vitro*. A) OV81 and OV81-CP40 and B) PEO-1 and PEO-C4.2 cells were treated with increasing doses of Olaparib alone or in combination with SMAP-061, and SMAP-061 alone or combination of both (C) OV81 and OV81-CP40; and D) PEO-1 and PEO-C4.2), to measure cell viability at 24 hours using MTT. Data presented as the mean ± SD (n=3). E) Combination index (CI) was calculated using the proliferation combinatorial assays conducted in A) for all four cell lines. CI > 1: antagonistic; CI = 1: additive; CI < 1: synergistic. <u>Formula:</u> (EC_50_combination_/EC_50_SMAP-061_) + (EC_50_combination_/EC_50_PARPi_). F) Clonogenic assay of OV81, PEO-1 and PEO-C4.2 was performed and quantified for cells treated with 6µM SMAP-061, 100nM Olaparib and the combination of both, after 2 weeks. Data presented as the mean ± SD (n=3), (unpaired Student T-tests, comparing each treatment group relative to each other, *p < 0.05, **p < 0.01, ***p < 0.001, ****p < 0.0001). Raw data represented in Fig. S4A. G) Western blot analysis assessing cell death markers (cleaved Parp and cleaved Caspase 3) and DNA damage (γH2Ax) expression to further evaluate dose and H) time dependency profiles in OV81, PEO-1 and PEO-C4.2 cells treated with SMAP-061, Olaparib or combo.

### SMAP-061 potentiates PARPi-induced cell death by decreasing RAD51 and preventing DNA repair

Our previous results have shown that the chemical modulation of PP2A using SMAP-061 leads to the significant downregulation of multiple DDR, and specifically, RAD51, ultimately triggering synthetic lethality (Fig. 3B-D). Furthermore, we show that SMAP-061 can potentiate the effects of PARPi in cells with varying HR status and sensitivities to platinum (Fig. 4). However, the mechanism driving such outcomes is not fully clear. Thus, we decided to evaluate the effect of different doses and treatment schedules of SMAP-061 alone and in combination with Olaparib, on the HR and cell cycle proteins previously shown to be altered by SMAP-061 treatment alone. Western blotting analysis showed that DNA damage signaling via increased phosphorylation of the sensor proteins RPA (Fig. 5A and Fig. S5A-S5F) and H2Ax (Fig. 4G and Fig. 4H) was significantly increased in the SMAP-061 only group, and in a dose dependent manner. The combinatorial effects on p-RPA and γH2Ax was most significant for OV81 and PEO-C4.2, as these cell lines were more resistant to PARPi alone due to no HR deficiency. Thus, SMAP-061 treatment was able to sensitize these two cell lines to Olaparib and lead to a more significant induction of p-RPA and γH2Ax, than either treatment alone. Nevertheless, because PEO-1 harbors a BRCA2 mutation, baseline activation of DNA damage with PARPi alone was significantly higher, showing little to no combinatorial potential in inducing these markers when combined with SMAP-061 (Fig. 5A). A decrease in protein expression of BRCA1/2, RAD51 (Fig. S5G-S5O) and the cell cycle regulators Rb, WEE1 and Securin (Fig. S5P-S5U) was also observed. As expected, and previously seen by SMAP-061 alone, the induction of other transducers of the HR pathway (Fig. S5S-S5U) was also achieved by the combination treatment. Interestingly, comparing SMAP-061 alone with SMAP + Olaparib treatment (Fig. 5A, lanes 2-4 vs 6-8; 10-12 vs 14-16; and 18-20 vs 22-24), we showed that the protein levels of RAD51 were substantially decreased in the combo treatment groups, even at the combination of Olaparib with the lowest dose of SMAP-061, in all 3 cell lines (Fig. 5A and Fig. S5G-S5I). Moreover, this data also suggests that SMAP-061 can exert equally efficient tumor suppressive and anti-proliferative effects when used at lower doses and in combination with Olaparib (lower EC_50_ – Table S3B), to when it is used at higher doses as a single agent. Other transducers and effector targets of the DDR pathway were also validated using western blotting analysis and presented similar expression patterns (Fig. S5S-S5U and Fig. S5J-S5O). These results further support the global additive effects of SMAP + Olaparib combination on engaging multiple transducers and effectors downstream of the DDR and HR pathways (Fig. S5A-S5U). Previous studies report that the clinically achievable concentration of Olaparib is ∼100µM(47,48). We further confirmed Olaparib at this dose showed a slight increase in DNA damage after 24 hours. However, SMAP-061 was more potent and the two drugs combined show a more robust effect (Fig. S5V) Furthermore, increased apoptosis was also observed when the two drugs were combined compared to each alone (Fig. S5V). Therefore, this data suggests that DNA damage and apoptosis were still efficiently induced at longer time points when using clinically achievable concentrations, further supporting our previous observations.

**Figure 5.**
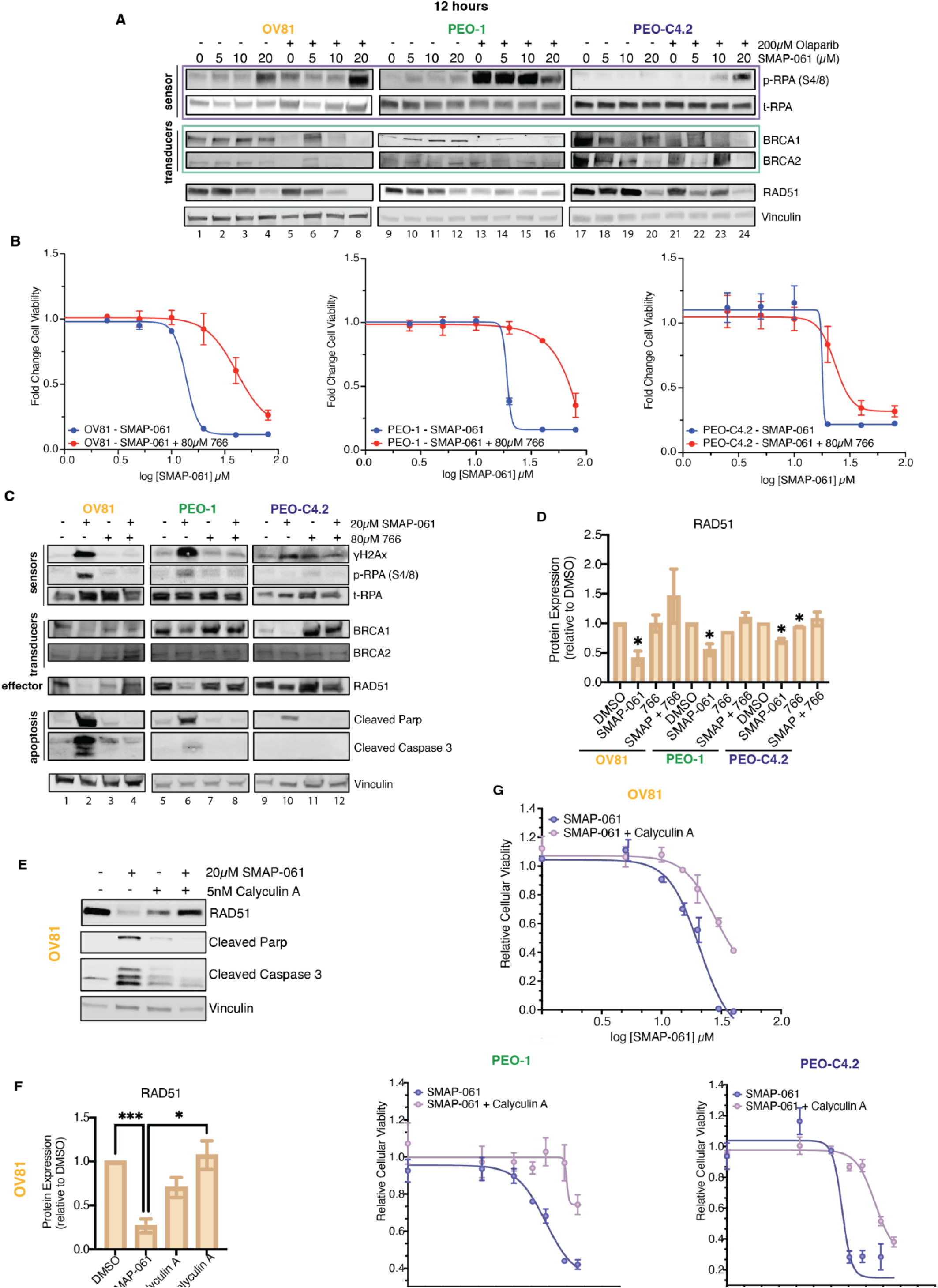
Combination of SMAP-061 and PARPi synergistically engages the DDR pathway and prevents DNA repair by specifically targeting and downregulating RAD51. A) Western blot analysis evaluating dose dependency effects of SMAP-061, Olaparib or combination on DDR protein expression and activity. **Sensor, transducer**, and **effector** proteins (from Fig. 3A schematic) were analyzed in OV81, PEO-1 and PEO-C42. B) OV81, PEO-1 and PEO-C4.2 cell lines were exposed to increasing doses of either SMAP-061 alone (blue) or in combination with 80µM of 766 (a biologically inactive analogue of SMAP-061) (red) and cell viability was measured at 24 hours by MTT analysis, to generate an EC50 curve. Data is presented as the mean ± SD (n=3). C) Rescue experiments were performed on OV81, PEO-1 and PEO-C42 cells treated with DMSO, SMAP-061, 766 or SMAP + 766. Rescue profile analysis of DDR (BRCA1/2 and RAD51) protein expression as well as recovery in cell death markers (cleaved Parp and cleaved Caspase 3) and DNA damage signaling capacity (γH2Ax and p-RPA) after 12 hours of treatment is analyzed by western blotting. Values underneath RAD51 blot represent the fold change of protein expression to its DMSO control sample, after Vinculin normalization. D) Quantification of RAD51 protein expression, with data presented as the mean ± SEM (n=3), (unpaired Student T-tests, comparing each treatment group relative to its respective DMSO, *p < 0.05). E) Calyculin A (PP2A Catalytic subunit inhibitor) rescue experiments were performed using Western Blotting (and respective quantification in F)) and G) MTT techniques to assess cellular viability and EC50 shift for OV81, PEO-1 and PEO-C4.2. 5nM of Calyculin A was preincubated for 1 hour, followed by SMAP-061 treatment. Data presented as the mean ± SD (n=3), (unpaired Student T-tests, comparing each treatment group relative to each other, *p < 0.05, ***p < 0.001).

Next, we sought to confirm that the effects of SMAP-061 on DDR proteins and induction of apoptosis is dependent on PP2A activation. Do to so, we conducted competition studies using a biologically inactive SMAP analog, 766(49,50). We have previously shown that compound 766 has similar chemical structure and is known to bind the same pocket in the PP2A holoenzyme as our active analogs, yet this molecule does not stabilize or activate the trimeric form of PP2A, thus resulting in no changes in cell viability. This tool compound represents a highly specific control for the observed changes in target protein expression given its high structural similarity to SMAP-061 and its ability to bind but not activate the phosphatase. Therefore, we treated cells with either DMSO (control), SMAP-061, 766, or the combination of SMAP-061 + 766 and analyzed essential molecular targets from the DDR and cell cycle pathways. To ensure successful competition and reversibility of PP2A’s activity and stability, we used 4 times the concentration of 766 as the concentration for the active SMAP-061, which was able to significantly compete off for the phosphatase binding and therefore reverse cell death profiles (Fig. 5B). Interestingly, the effects of SMAP-061 on DNA damage sensing mechanisms (γH2Ax and p-RPA expression) were fully reversed in cells treated with SMAP-061 and 766 combinations, after 12 hours (Fig. 5C lanes 2 vs 4; 6 vs 8; and 10 vs 12, and Fig. S6A-S6C). Additionally, the protein expression levels of both the G_1_ cell cycle checkpoint protein Cyclin D1 and the cell death markers, cleaved PARP, and cleaved Caspase 3, were also reversed in the co-treated cells when compared to SMAP-061 treatment alone (Fig. 5C, Fig. 5D and Fig. S6D). However, BRCA1 and BRCA2 expression was not rescued in all cellular models (Fig. 5C, Fig. S6E and Fig. S6F), indicating these proteins are not the main driving factor of the SMAP-061 mediated apoptosis. Notably, the RAD51 protein loss was completely reversed, suggesting this target is necessary for SMAP-061 mediated cell death. In support of the latter statement, comparing RAD51 expression intensity in SMAP-061 only conditions (Fig. 5C, lanes 2, 6 and 10), showed that OV81 and PEO-1 cells had a more dramatic decrease in RAD51 when compared to PEO-C4.2 (Fig. 5D). Therefore, the degree of RAD51 protein expression was significantly and inversely correlated with the expression levels of γH2Ax, p-RPA, cleaved PARP and cleaved Caspase 3, for all three cell lines, as confirmed in our correlation studies (Fig. S6G).

To further confirm the specificity of the SMAP-061 modulating PP2A’s activity we also used Calyculin A – a highly specific PP2A and PP1 Catalytic subunit inhibitor(51). Using similar rescue experiments as we did for 766 (Fig. 5B), we were able to demonstrate that SMAP-061-mediated effects directly depend on PP2A modulation to induce the previously observed phenotypes (Fig. 5E and Fig. 5F). Our cell viability assays show that Calyculin A shifts the EC_50_ of SMAP-061 when preincubated for an hour, priming PP2A to remain inactive despite of SMAP-061 presence. At the highest SMAP-061 dose, 40µM, Calyculin A can rescue cell death by 40%, comparatively to SMAP-061 treatment alone, where 100% of cells are no longer viable (Fig. 5G). Using western blotting techniques, we also evaluated the specificity of SMAP-061 in modulating PP2A’s activity within the HR and cellular death molecular pathways (Fig. 5E). We observed that for all three cell line models, SMAP-061 action is effectively prevented by Calyculin A preincubation when used in combination, as observed by the cell death markers as well as RAD51 protein expression rescue.

In sum, this data does provide evidence that SMAP-061 works through the activation of PP2A and the downregulation of RAD51 may be directly involved with synthetic lethality effects seen in both HR-deficient and proficient HGSC tumors treated with SMAP-061 alone or in combination with PARPi.

### SMAP-061 improves *in vivo* survival and significantly decreases tumor progression in PDX mouse models, as a single agent and in combination with PARPi

So far, we have established that PP2A activation leads to apoptotic effects *in vitro*, through targeting essential DDR, HR, and cell cycle proteins, which results in an irreversible ‘BRCAness’ phenotype. We also showed that, in combination with PARPi, SMAP-061’s anticancer activity is enhanced, triggering synthetic lethality in HGSC cells. Given our recent discoveries and the promise these compounds hold clinically, we evaluated their efficacy *in vivo*, both alone and in combination with Olaparib. Three independent PDX tumor models were utilized, including OV81, OV262 and OV17 (Table S2C), where we conducted efficacy studies to evaluate tumor growth and survival. We found that in the OV81 model Olaparib alone was not effective in improving survival (Fig. 6A) or decreasing tumor burden (Fig. 6B), as this PDX was obtained from the ascites of a women diagnosed with HGSC who did not harbor a HR-deficient mutation. However, SMAP-061 treatment alone and in combination with Olaparib showed significant improvement in median survival (Fig. 6A). This increase in survival was consistent with the significant tumor growth inhibition seen in both the SMAP-061 alone and combination groups which was maintained until the end of the study (Fig. 6B). Interestingly, the synergistic effects observed in our *in vitro* models were not replicated in our *in vivo* studies which indicate a need to modify the dosing and schedule regimen. Furthermore, γH2Ax staining showed that SMAP-061 + Olaparib treatment group did have higher intensity (Fig. 6C and Fig. S7D). In the OV262 PDX model, a HGSC tumor obtained from a patient that harbored a germline BRCA2 mutation (p.N3124I), we found that SMAP-061 alone was as efficacious in inhibiting tumor growth as the single agent Olaparib (Fig. 6D and Fig. 6E). However, the combination of these two agents resulted in significant tumor regression, which was not achieved by either one of these compounds alone. These results highlight that the SMAP-061 + Olaparib combination therapy strategy has synergistic effects in reducing tumor burden in HR-deficient HGSC tumor models. γH2Ax staining reveals that neither SMAP-061 or Olaparib alone can significantly increase foci formation, however the combination of the two drugs significantly increases its accumulation (Fig. 6F and Fig. S7E). The lack of γH2Ax staining in tumors treated with single agents could be due to these cells being proficient in resolving DNA damage prior to exacerbated accumulation. However, in the combination treatment groups, the tumor cells do not overcome the chronically accumulated DNA insults, thus resulting in tumor regression. Finally, we used another HRD PDX mouse model, OV17, which harbors a BRCA1 germline mutation <u>(p.R762fs</u>). Our data shows that SMAP-061 alone had higher efficacy in affecting tumor weight (Fig. 6G) and % tumor change (Fig. 6H) when compared to the single agent Olaparib. Importantly, despite significant tumor growth inhibition in both single agent groups, SMAP-061 + Olaparib combination induced tumor regression (Fig. 6H). Histology shows that γH2Ax foci are significantly higher in OV17 PDX tumors treated with SMAP-061 alone or SMAP-061 + Olaparib combination (Fig. 6I and Fig. S7F). In all the treatment studies there was no significant weight loss compared to the Vehicle control treated animals (Fig. S7A-S7C), as previously shown in other *in vivo* SMAP-061 studies.

**Figure 6.**
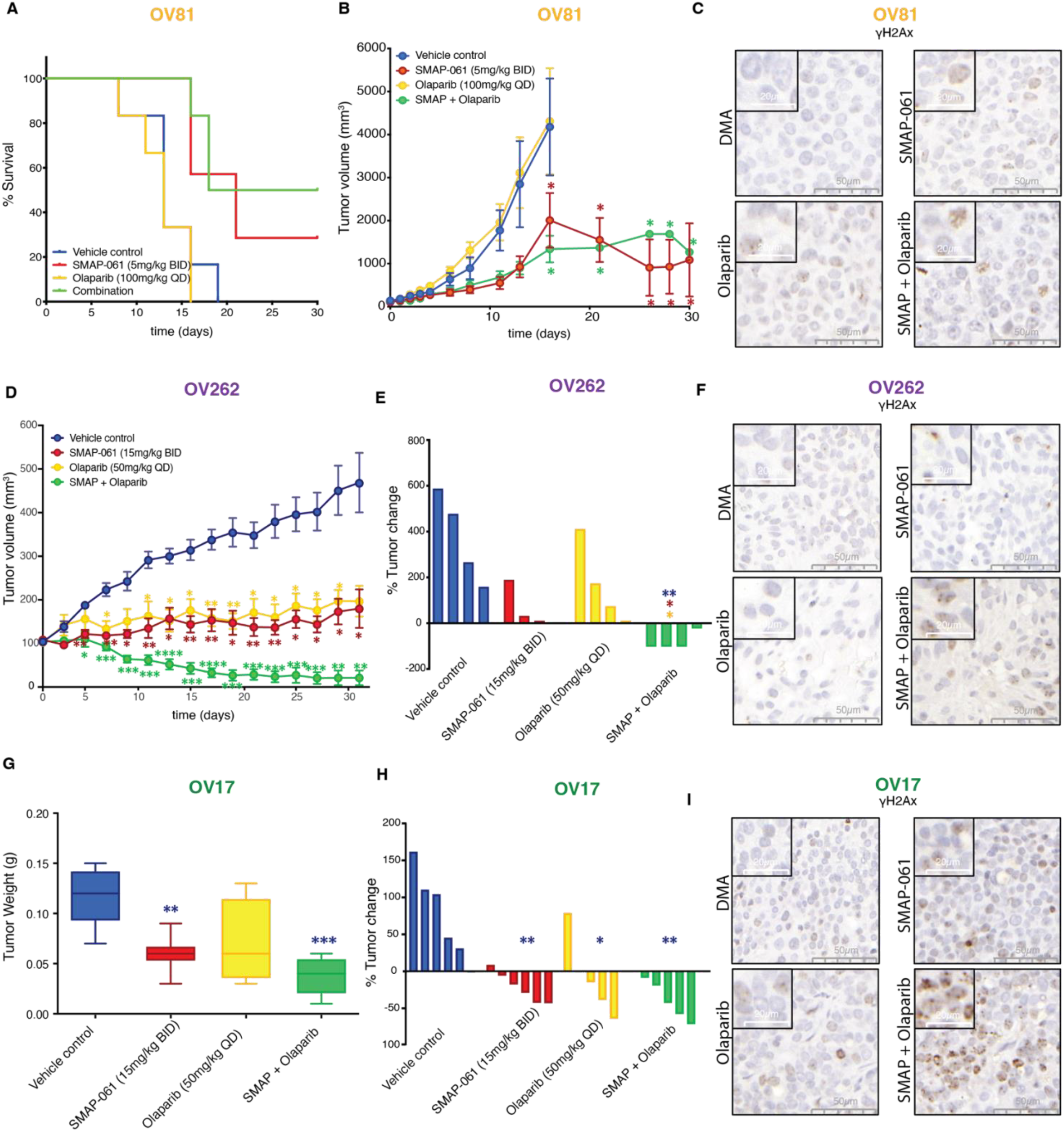
Effects of SMAP-061 *in vivo* show improved survival and significant tumor regression, as a single agent and in combination with PARPi, in both BRCA1/2 wild-type and mutant HGSC PDX tumors. OV81 PDX studies were conducted with tumors implanted in the right flank of NSG mice and allowed to grow between ∼100-200mm^3^ before enrollment in one of 4 treatment groups: Vehicle control (n=6), 5mg/kg SMAP-061 (n=6), 100mg/kg Olaparib (n=6) or SMAP + Olaparib combination (n=6). Tumors were measured every other day. Data plotted as a function of A) mouse survival and B) tumor volume (efficacy) over time. Survival threshold was selected to be 1000mm^3^ (tumors < 1000mm^3^: survival; tumors > 1000mm^3^: no survival). Data presented as mean ± SEM (Student T-tests, comparing each treatment group relative to vehicle control, *p < 0.05). C) Histology slides with tumor samples stained for γH2Ax (brown), representative of each individual treatment group in the OV81 PDX *in vivo* study. 50µm and 20µm scales are included for each respective picture. D) Efficacy study of OV262 PDX tumors, implanted in the right flank of NSG mice and allowed to grow between ∼90-120mm^3^ before enrollment in one of 4 treatment groups: Vehicle control (n=4), 15mg/kg SMAP-061 (n=3), 50mg/kg Olaparib (n=4) or SMAP + Olaparib combination (n=4). Tumors were measured every other day. Tumor volume was calculated and plotted over time (31 days). Data presented as mean ± SEM (Student T-tests, comparing each treatment group relative to vehicle control, *p < 0.05, **p < 0.01, ***p < 0.001, ****p < 0.0001). E) Waterfall plot of OV262 tumor volume change (from day 0 to day 31) comparing all treatment groups to each other. Data presented as mean ± SEM (unpaired Student T-tests, comparing each individual treatment group relative to each other, *p < 0.05, **p < 0.01). F) Histology slides with tumor samples stained for γH2Ax (brown), representative of each individual treatment group in the OV262 PDX *in vivo* study. 50µm and 20µm scales are included for each respective picture. G) and H) OV17 PDX studies were conducted with tumors implanted in the right flank of NSG mice and allowed to grow between ∼80-250mm^3^ before enrollment in one of 4 treatment groups: Vehicle control (n=6), 15mg/kg SMAP-061 (n=6), 50mg/kg Olaparib (n=5) or SMAP + Olaparib combination (n=5). Tumors were measured every other day. Data plotted as a function of G) tumor weight post sacrifice and H) tumor change percentage (waterfall plot) for each treatment group. Data presented as mean ± SEM (unpaired Student T-tests, comparing each individual treatment group relative to each other, *p < 0.05, **p < 0.01, ***p < 0.001). I) Histology slides with tumor samples stained for γH2Ax (brown), representative of each individual treatment group in the OV17 PDX *in vivo* study. 50µm and 20µm scales are included for each respective picture.

Together, these results further support that SMAP-061 can effectively inhibit tumor growth in both BRCA1/2 mutant and wt tumors *in vitro* and *in vivo*. Thus, when combined with PARP inhibitors, SMAP-061 significantly induces tumor regression in BRCA1/2 mutant tumors. Importantly, this data highlights the therapeutic potential of SMAP-061 for the treatment of HGSC and points to the possibility of PP2A modulation in inducing synthetic lethality, which could allow all patients to benefit from PARP inhibitor therapy regardless of HR status.

## Discussion

HGSC is a heterogenous disease driven by a high degree of genomic instability due to defects in HR pathway, for which genetic drivers that lead to its recurrence and drug resistance have not yet been fully established. Even though the discovery of the BRCA genes revolutionized the field for both ovarian cancer screening and prevention, allowing PARPi to thrive clinically(6– 8), only 21% of HGSC patients harbor germline (15%) or somatic (6%) BRCA1/2 mutations(52), resulting in a distinct subset of HGSC patients that are able to benefit from these therapies. Therefore, it is essential to identify additional unique and targetable genetic drivers of HGSC and introduce novel targeted therapeutic opportunities that take advantage of HGSC tumors dependency on the HR pathway.

The PP2A family of genes has been extensively studied in cancer and it has been recognized as a major tumor suppressor and main negative regulator of a series of oncogenic pathways. Its Serine/Threonine phosphatase activity drives the dephosphorylation of a vast array of downstream substrates and ultimately regulates cellular homeostasis. While targeted reduction of PP2A has been shown to be essential for the transformation of the non-malignant human FTSEC model(34), the genetic and molecular impact each PP2A family member may have in HGSC tumorigenesis and disease progression has not been established. Unlike most tumor suppressors, such as p53 and others, which are found to be functionally inactivated in cancer via genetic mutations, PP2A heterotrimeric components are rarely mutated in human cancers. PP2A inactivation in cancer primarily involves non-genetic mechanisms, including overexpression of endogenous inhibitors or suppressive post-translational modifications to the C terminal tail of its catalytic domain(15). As noted here in this current manuscript, we uncovered that >90% of HGSC tumors harbor heterozygous loss and decreased mRNA expression of PP2A genes important to malignant transformation and/or holoenzymatic core formation, further correlating with decreased overall patient survival. Thus, PP2A tumor suppressor activation represents an attractive therapeutic alternative for the treatment of HGSC. Nevertheless, protein phosphatases have been largely understudied as a target for drug development, mostly due to their perceived ‘undruggable’ nature. However, in this current study, we uncover that SMAP-061 can potently induce apoptosis in numerous PD HGSC models and potentiate PARPi mediated cell death independent of platinum sensitivity and HR status. SMAP-061 belongs to a novel class of small molecules that have shown to directly bind to and promote the formation and stabilization of PP2A tumor suppressive holoenzymes which ultimately modulates PP2A activity(27)^-51^. Mechanistic studies have shown that SMAP-061 selectively binds PP2A-Aα and -Cα, and recruits PP2A-B56α subunit for a trimeric holoenzyme formation, as determined through Cryo-EM structures(26). Upon binding, SMAP-061 is shown to further increase PP2A-Cα subunit carboxymethylation, which correlates with enhanced enzymatic activity of the phosphatase, therefore rendering its most active form. Furthermore, disruption of this binding pocket confers cellular resistance to SMAP-061, suggesting that these molecules trigger cancer cell death and impede tumor growth(28,53,54). These studies show that tumors with mutations in the A scaffolding subunit (K194R and L198V) located at the SMAP-061 binding site interface displayed resistance to treatment *in vivo*. Interestingly, alternative mechanisms have also been proposed wherein SMAP-061 preferentially stabilizes the B55α-PP2A heterotrimer in blood cancers(30). These studies were performed through a CRISPR-Cas9 functional screen where each of the subunits were knocked out (KO) and enzymatic activity was measured. Pulldown experiments further showed that that B55α specifically binds to PP2A-Cα under SMAP-061 exposure, utilizing leukemic models. It is not fully understood why there are discrepancies between these studies. However, cell-specific differences can contribute to these disparities given the impact of the abundance and/or post-translational modifications of B55α and B56α subunits and solid tumor models from mouse xenografts vs *in vitro* suspension cell line models is important. PP2A members of the B56 family have been shown to play essential roles in cell adhesion and epithelial cell junctions, as PP2A-B56 inhibition is associated with desmosomal disruption and motility regulation(55,56). Additionally, CRISPR KO screen approaches for the PP2A regulatory genes induce compensatory upregulation of other B subunits with overlapping downstream targets, thus requiring careful interpretation of the observed phenotypic results(30,57). Nevertheless, despite slight contradiction among studies, multiple groups consistently agree that studying these SMAP molecules is essential to further evaluate the therapeutic potential of PP2A modulation, and further explore the mechanisms by which they exert their potent anticancer properties with limited tolerability concerns, even in long-term dosing studies *in vivo*(26–30). In our current study, we provide experimental evidence that SMAP-061 is highly selective and specific in modulating PP2A’s activity and confirm that the effects of SMAP-061 on DDR proteins and induction of apoptosis is dependent on PP2A activation by using two different methods. The first is through the use of a biologically inactive SMAP analog, 766, which we have previously shown has similar chemical structure and can bind the same pocket in the PP2A holoenzyme as our active analogs, yet does not stabilize or activate the trimeric form of PP2A, thus resulting in no changes in cell viability^48,50^. Using this compound, we were able to show that SMAP-061 induced DNA damage and apoptosis were abrogated and the degradation of RAD51 protein was eliminated suggesting this target is necessary for SMAP-061 activity and mediated cell death. Secondly, to further confirm the specificity of the SMAP-061 in activity PP2A and the role of RAD51 in regulating its impact on cell death, we utilized Calyculin A, a highly specific PP2A and PP1 Catalytic subunit inhibitor(51). We found that SMAP-061 induced degradation of RAD51 is effectively prevented by Calyculin A preincubation and apoptosis is abrogated. In sum, this data does provide evidence that SMAP-061 works through the activation of PP2A; however further studies are still required to better understand which regulatory subunit(s) of PP2A are driving these anticancer effects, to better leverage their therapeutic potential.

As more than 70% of HGSC tumors are characterized by HR defects, RAD51 plays a critical role in the tumorigenesis of these cancers, by being an essential component of HR that modulates DNA strand exchange in DSB repair. The inhibition of RAD51 has been shown to lead to synthetic lethality through the accumulation of γH2Ax, thus spearheading the clinical development of RAD51 inhibitors^57,59–61^. By showing that SMAP-061 can potently reduce RAD51 leading to concomitant apoptosis, our studies introduce a unique mechanism of synthetic lethality, through a small molecule mediated modulation of PP2A (Fig. 7A). These results suggest that SMAPs could be effective against tumors that HR-proficient as well as PARPi resistant tumors given that RAD51 upregulation is a common mechanism of resistance. The mechanism by which SMAP-061 is leading to RAD51 decrease of expression remains unclear; however given that the loss of the B55*α* regulatory subunit (PPP2R2A) has been shown to inhibit HR DNA repair and predict tumor sensitivity to PARP inhibition via the downregulation of RAD51(24), SMAP-061-mediated displacement of B55*α* and recruitment of PP2A-B56*α* could be affecting RAD51 stability. Even though PP2A and RAD51 direct interaction has never been established and further studies are warranted to assess such hypothesis, published evidence has clearly demonstrated that PP2A plays an indirect role on RAD51’s activity and protein stability. The BRCA2–PP2A-B56 complex is essential for the efficient RAD51 filament formation at the sites of DNA damage(25). Furthermore, PLK1, which has been show to directly phosphorylate and activate RAD51’s recruitment(61), is dephosphorylated and inactivated by PP2A(62,63), reinforcing the idea that PP2A plays an indirect role on RAD51’s activity. Further studies are necessary to assess the mechanism by which PP2A is regulating RAD51 and determine if putative phospho sites on RAD51 are dephosphorylated by PP2A, leading to the initiation of RAD51 protein degradation cascades, thus preventing a fully efficient HR repair mechanism to be properly engaged (Fig. 7A).

**Figure 7.**
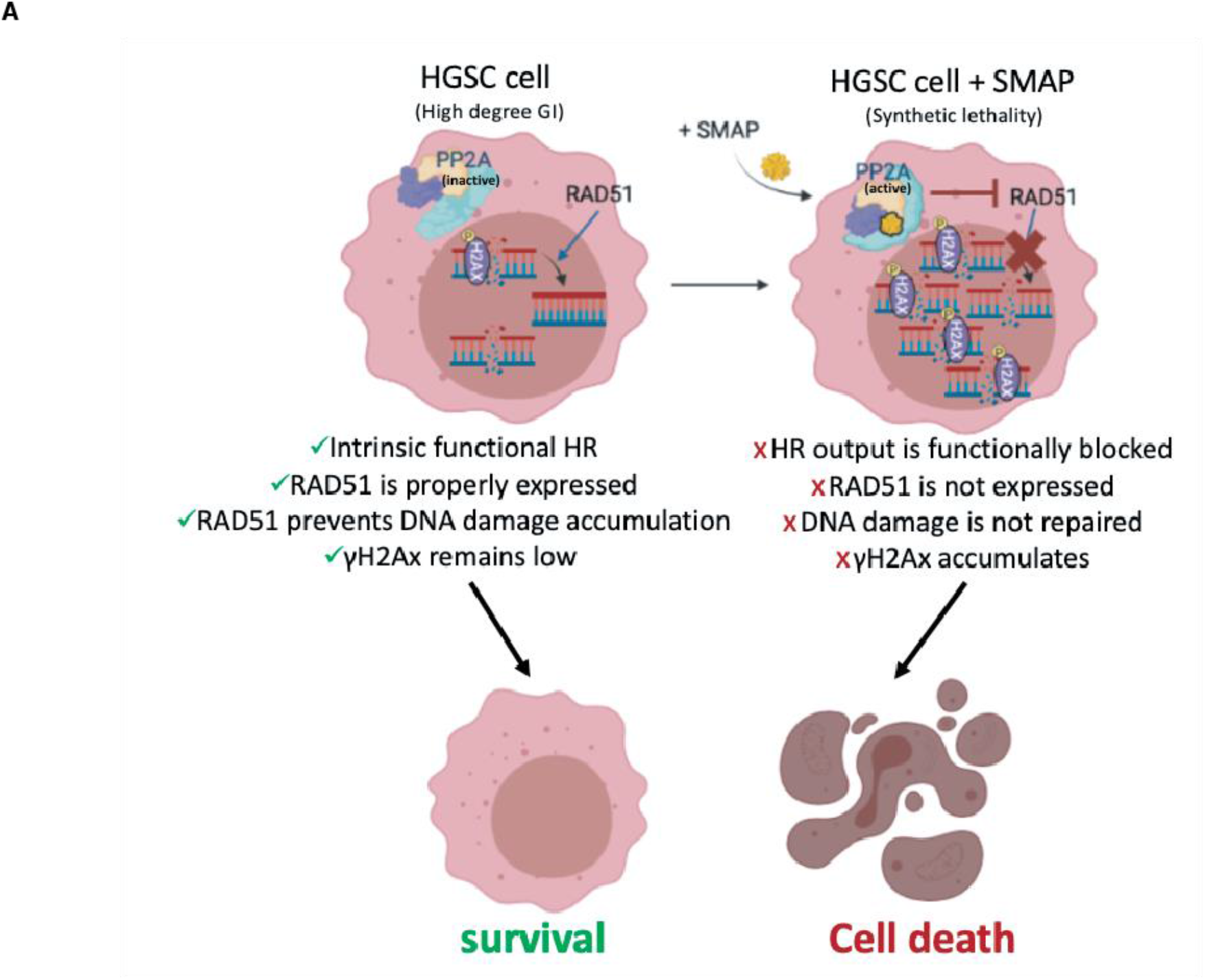
RAD51 inhibition leads to the chronic accumulation of γH2Ax and DNA errors inherent from genomic instability, an ovarian cancer trait. A) HGSC cells harbor a high degree of Genomic Instability (GI) which results in small nicks and breaks in the DNA over time. Sensor proteins such as γH2Ax are recruited to the site of damage to prevent DNA degradation and remain in check until a repair mechanism is triggered. The minimally functional HR machinery is activated and prompts the recruitment of the main downstream effector protein, RAD51, which by keeping those damaging signals at bay, allows cancer cells to survive, even if with relatively higher baseline levels of γH2Ax when compared to normal cells. For this reason, ovarian cancer cells are known to functionally tolerate and survive high baseline levels of DNA damage while avoiding apoptotic cues (left). When SMAP-061 is added to the cell, PP2A’s anti-tumor activity is functionally restored, and synthetic lethality is induced (right). RAD51 expression is inhibited by PP2A, preventing the HR output to be efficiently propagated. This results in the incapacity of the cells to repair their inherent DNA damage, thereby leading to the accumulation of γH2Ax signals in the nucleus, and ultimately, triggering apoptosis.

Furthermore, the functional implications of the high percentage of HGSC tumors that harbor loss of PP2A genes it remains unclear. For example, the effect of PP2A’s loss on DDR during HGSC development and response to PARPi needs to be further explored and studies need to be conducted regarding the role of RAD51 in the SMAP-dependent cellular effects seen in HGSC. RAD51 is essential to ensure the success of HR in repairing and resolving DNA damage; and without it, the homology search and strand invasion steps unique to HR cannot be successfully executed. Unlike RAD51 and other HR regulators, BRCA1 and BRCA2 mutation carriers are significantly more prevalent, and pathogenic variants of these genes as well as their functional output have been best studied in HGSC and breast cancer. Mutations in BRCA1 and BRCA2 genes are functionally relevant for tumor development and resistance and ultimately lead to HRD, as a result of the inability of these cells to form RAD51 filaments, resulting in a failure to accomplish homology search and strand invasion of the sister chromatid(60). Thus, targeting proteins downstream the BRCA genes in the HR pathway, more specifically RAD51, seems to represent a more beneficial and advantageous strategy to tackle clinical challenges and overcome PARPi resistance driven by BRCA1/2 mutation reversions. Targeting RAD51 has proven biologically efficacious for the inhibition of human cancer progression(49,58–60,64,65). Data shows that RAD51 antisense oligonucleotides and RNA interference as well as specific small molecule inhibitors that reduce RAD51 expression, re-sensitized multiple cancer types to chemo and radiation therapies, proving RAD51 to be a suitable target for therapy. Despite the lack of viable potent, direct, and highly specific inhibitors of RAD51, thus these PP2A modulators seem to be highly effective in regulating RAD51 expression and/or stability, presenting a great promise to be used in the clinic as an indirect RAD51 inhibitor. Beyond HGSC, these studies also lay the foundation for PP2A modulators for the treatment of other cancers that may harbor BRCA1 and BRCA2 mutations or RAD51 dependency given that the two PDX models used in these studies harbor BRCA1 and BRCA2 mutations that are also commonly seen in breast cancers(50,66). Distinct from HGSC, breast cancer patients that harbor BRCA1 or BRCA2 mutations have not shown as promising response to PARP inhibitor therapy therefore PP2A modulation alone or in combination with PARPi may represent a more effective therapeutic approach(67).

In summary, we have established a novel mechanism by which SMAP-061 target and induce programmed cell death in HGSC cells and have further elucidated a new role of PP2A in regulating DDR in HGSC (Fig. 7A). Moreover, a deeper molecular and mechanistic understanding of the antitumor effects of SMAP-061 has allowed us to introduce novel pharmacodynamic markers of PP2A target engagement and therapeutic efficacy, such as RAD51. More importantly, we believe that introducing SMAP-061 as a novel targeted therapy for the treatment of HGSC can further optimize treatment selection and patient stratification. Such advancement can help bring tremendous advances to the field and further leverage drug development and discovery for the treatment of HGSC, while also preventing treatment resistance from developing.

## Materials and Methods

### Generation of patient cell lines and PDX models

Patient samples were collected under an approved University Hospitals of Cleveland Institutional Review Board (IRB). The IRB protocol included the prospective collection of discarded tissue and the generation of PDX with written informed consent obtained from the study subjects (PI: DiFeo). As part of these IRB protocols, clinical and pathological data were gathered for some patients, and included age at diagnosis, race, tumor stage and grade, histological type, and overall survival. The PD cell lines (OV78, OV81, OV82, OV84, OV85, OV93, OV145 and OV262) were generated from primary human ovarian cancer tissue following protocols previously described(68). To generate PDX models, tumors or ascites were removed from patients and ∼2 mm^3^ tumor implants were grafted, or cells were injected subcutaneously in NSG mice, respectively. Once tumors were detected, tumor volumes were measured weekly and were collected after reaching ∼1000 mm^3^. Tumors were then (a) re-implanted into another NSG mouse as a passaging tool, (b) processed using IHC analysis to compare the pathology to the original patient tumor, (c) processed for exome sequencing, and lastly (d) frozen in freezing media for future drug studies.

### *In vivo* drug studies

Mice were randomized to one of 4 different treatment groups depending on their initial tumor size. Mice were enrolled when tumors reached ∼100mm^3^ and distributed in different treatment groups so that the initial tumor volume average was the same across each group. Mouse tumor volumes and body weight were measured every other day. SMAP-061 was diluted in 10% DMA (Sigma Aldrich 271012-12), 10% Solutol (Sigma Aldrich 42966) and 80% warm RNA-free water. DMA control and SMAP-061 were administered via oral gavage twice a day while PBS control and Olaparib were injected via IP once a day.

### Proliferation and Colony Forming assays

HGSC cells were seeded in a 12-well plate overnight to reach 70% confluency at the day of treatment. Cells were then treated with either DMSO (Thermo-Fisher Scientific BP231-100), SMAP (5, 10, 20, 30 or 40µM), cisplatin (5, 10, 20, 40 or 80µM) or Olaparib (25, 50, 100, 200, 600µM) (Selleck Chemicals S1060) and incubated for 24 hours. Cell viability was measured by MTT assay using a 3-(4,5-dimethylthiazol-2-yl)-2,5-diphenyltetrazolium bromide kit (Thermo-Fisher Scientific M6494). For the isobologram assays, the combination of dose increments of SMAP and Olaparib was utilized, and cell viability was measured after 24 hours of incubation with the drugs. To perform colony forming assays, cells were plated at a low density (∼100 cells/well) in biological triplicate in a 6-well plate. After 24 hours, cells were treated with DMSO, 6µM of SMAP, 100nM of Olaparib or 6µM SMAP + 100nM Olaparib and incubated for 10-12 days. Drug medium was replaced every three days. At day 12, cells were fixed and stained with 1% crystal violet solution. Image lab software was used to quantify colony forming density, measured by pixel intensity.

### Western Blotting

Cells were harvested and pelleted for protein extraction using a cocktail consisting of 1X RIPA Lysis and Extraction Buffer (EMD Millipore 20-188), 5% glycerol, protease inhibitor (ThermoFisher Scientific A32955) and phosphatase inhibitor (Thermo-Fisher Scientific A32957). Quantification of the isolated proteins was performed using a Pierce BCA Protein Assay kit (ThermoFisher Scientific 23227), run on a 12% or 4-20% gradient SDS–polyacrylamide electrophoresis gel (BioRad 4568045 and 4568095, respectively). Proteins were transferred onto a nitrocellulose membrane (Bio-Rad 1704159) using the quick semi-wet transfer Trans-Blot Turbo transferring machine. Membranes were blocked using 5% non-fat milk (ThermoFisher Scientific 50488785) made with 1% Tris-Buffered Saline Tween20 (TBST) buffer (AMRESCO 10791-792). Antibodies were purchased from (1) Cell Signaling: p-Rb (8516), t-RB (9309 and 9313), Cyclin D1 (2922), PLK1 (4513), p-Cyclin B1 (4133), t-Cyclin B1 (4135), Cyclin E (4129), γH2Ax (9718), t-RPA (2208), BRCA1 (14823), BRCA2 (10741), p-ATM (13050), t-ATM (92356), p-CHK1 (2348), t-CHK1 (2360), p-CHK2 (2197), t-CHK2 (6334), RAD51 (8875), CDC25A (3652), Wee1 (13084), GAPDH (5174), Cleaved PARP (9541), Cleaved Caspase 3 (9661), Vinculin (13901); (2) Bethyl Lab: p-RPA (A300-245A); (3) Santa Cruz: Vinculin (sc-73614), GAPDH (sc-47724) and (4) EMD Millipore BRCA2 (OP95).

### Cell Cycle experiments

For cell cycle analysis, cells were plated in 10cm cell culture dishes at a density of 2M cells per plate in biological triplicate (approximately 70% confluency) and treated the next day. Cells were then treated with 2mM of Thymidine (Sigma-Aldrich T1895) and incubated for 24 hours to allow block of the cells in the S phase of the cell cycle. Cells were then released from the Thymidine block by replenishing with regular media for 6.5 hours to allow cells to reengage in cell cycle progression. Following the 6.5 hours release, cells were then treated with either DMSO or 20µM SMAP and incubated for 6, 10 and 16 hours and subsequently harvested and fixed using 70% cold ethanol. PI (ThermoFisher Scientific F10797) staining was performed according to FxCycle PI/RNAse Staining Solution protocol available online (www.thermofisher.com) and cell cycle profiles were registered using the Bio-Rad Zed flow machine. Cell cycle graphs were finalized utilizing FlowJo 8.

### Immunofluorescence

OV81 cells were grown on glass coverslips for 24 hours followed by 12 hours of exposure to 20µM SMAP-061. Cells were then fixed using 4% paraformaldehyde, permeabilized with 0.1% TritonX-100, and blocked with 3% BSA diluted in 1X PBS. Cells were then incubated with γH2AX primary antibody (Cell Signaling, 9718) overnight at 4°C. The cells were washed with 1X PBS and incubated with fluorescent secondary antibody for 1 hour at room temperature. Coverslips were mounted on glass slides using mounting media with DAPI (Invitrogen, P36966). Images were acquired using a Leica DMI6000 inverted microscope and then analyzed using Image-J.

### Histology

Immunohistochemical staining was performed on the DAKO Autostainer (DAKO, Carpinteria, CA) using Rabbit-on-rodent HRP (BioCare Medical, Pachero, CA, RMR622) and diaminobenzadine (DAB) as the chromogen. Heat induced epitope retrieval (10 mM Tris/1mM EDTA pH9) was used prior to staining for both antibodies. De-paraffinized sections were then labeled with negative (no primary antibody) and positive controls (RAD51 Abcam, ab133534) and γH2Ax (Cell Signaling, 9718), which were stained in parallel with each set of slides studied. γH2ax foci was scored unbiasedly by a Pathologist and quantification was further verified using Image-J.

### Statistical methods

Statistical methods used are described in each respective figure legend. Fold change controls and normalization methods are also specified for each one of the analyses.

## Acknowledgments

The authors would like to thank several core facilities within the Case Comprehensive Cancer Center (NIH P30CA043703) including the Athymic Animal and Preclinical Therapeutics Core. The authors would like to thank the Tissue and Molecular Pathology at Rogel Cancer Center at the University of Michigan (NIH P30 CA04659229) as well as the Cancer Center’s pilot funds in support of this project. In addition, we would like to thank all the generous donors and Foundations who have supported the DiFeo lab and strive to improve the outcomes of ovarian cancer patients including Norma C. and Albert I. Geller, The Silver Family Foundation and Jacqueline E Bayley (JEB) Foundation. Further acknowledgments are extended to Dr. Toshiyasu Taniguchi for providing us with the PEO-1 and PEO-C4.2 isogenic cell lines.

## Author contributions

RAA, AA, GC, NP, JC, AG, PLJ, RG, MD, AS, CMO and DSM performed experiments and/or analyzed data. RAA, CMO, JZ, GN and AD provided scientific interpretations to data analysis. RAA made the figures. AJA, SW, KZ, CN, KR, and SS provided patient samples. DT and SS provided immunohistochemistry analysis and pathology scoring. RAA, GN and AD interpreted data, wrote, and revised the manuscript. AD supervised overall design and study interpretation. All authors discussed results and provided input on the manuscript.

