## Supplemental Figures for "Small molecule mediated stabilization of PP2A modulates the Homologous Recombination pathway and potentiates DNA damage-induced cell death"

### Supplementary Figure 1

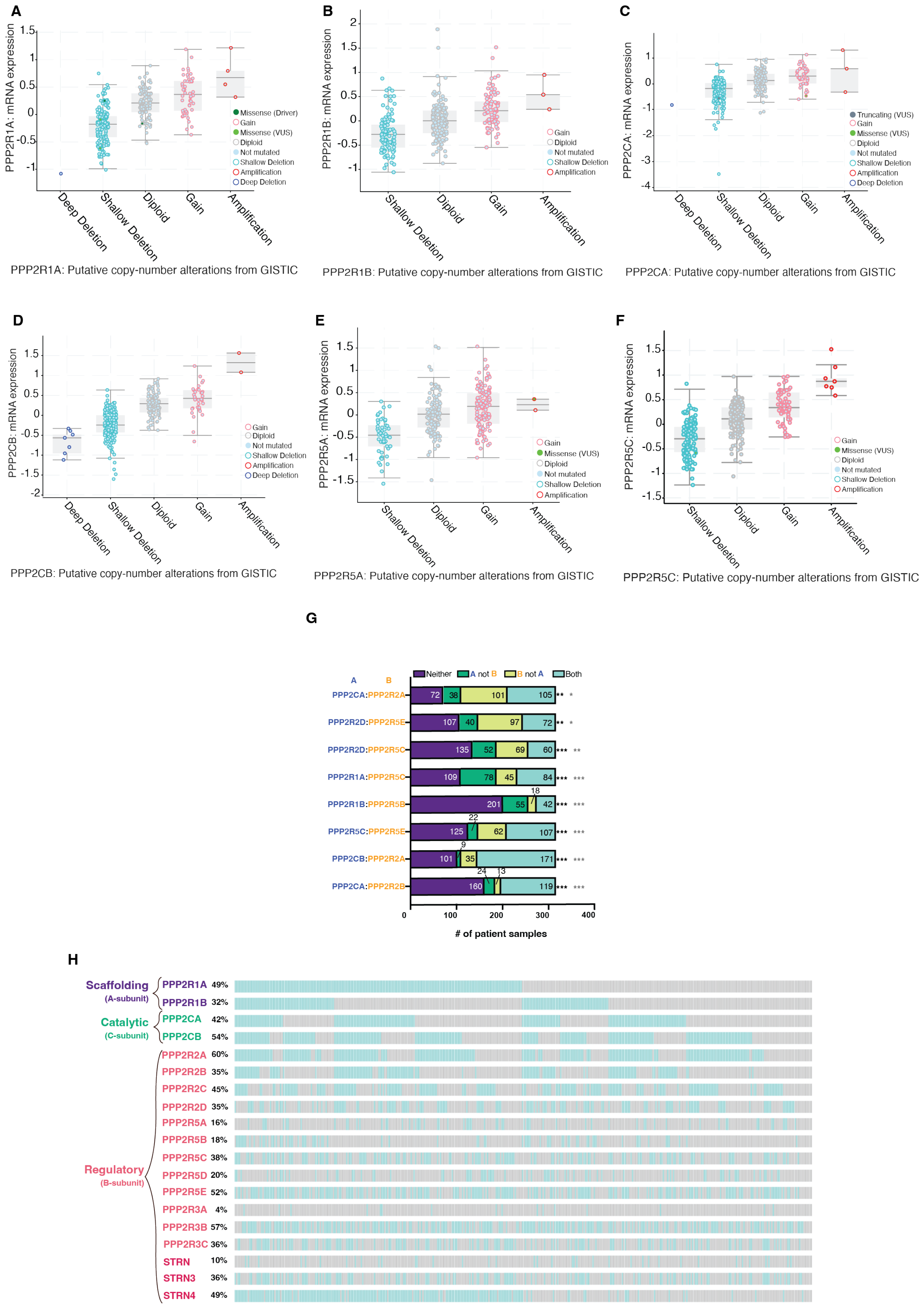

**Supplementary Figure 1 – PP2A family genes mRNA profiles show putative copy-number alterations proportionally correlate with their respective mRNA expression levels.** Shallow and deep deletion, diploid, gain and amplification copy-number alterations for A) PPP2R1A, B) PPP2R1B, C) PPP2CA, D) PPP2CB, E) PPPP2R5A, F) PPP2R5C, relative its own mRNA expression levels. G) Analysis of statistically significant co-occurrences in PP2A genes using panel C's data. ***p<0.001, **p<0.01; ***q<0.001, **q<0.01, *q<0.05. Tendency – co-occurrence. Statistical analysis provided by cBioPortal. H) Individual patient tumor data analysis showing frequency of PP2A genes heterozygous loss as well as co-occurrence profiles. Each bar represents an individual tumor, for which grey signifies negative and blue positive for Hetloss.

Supplementary Figure 2

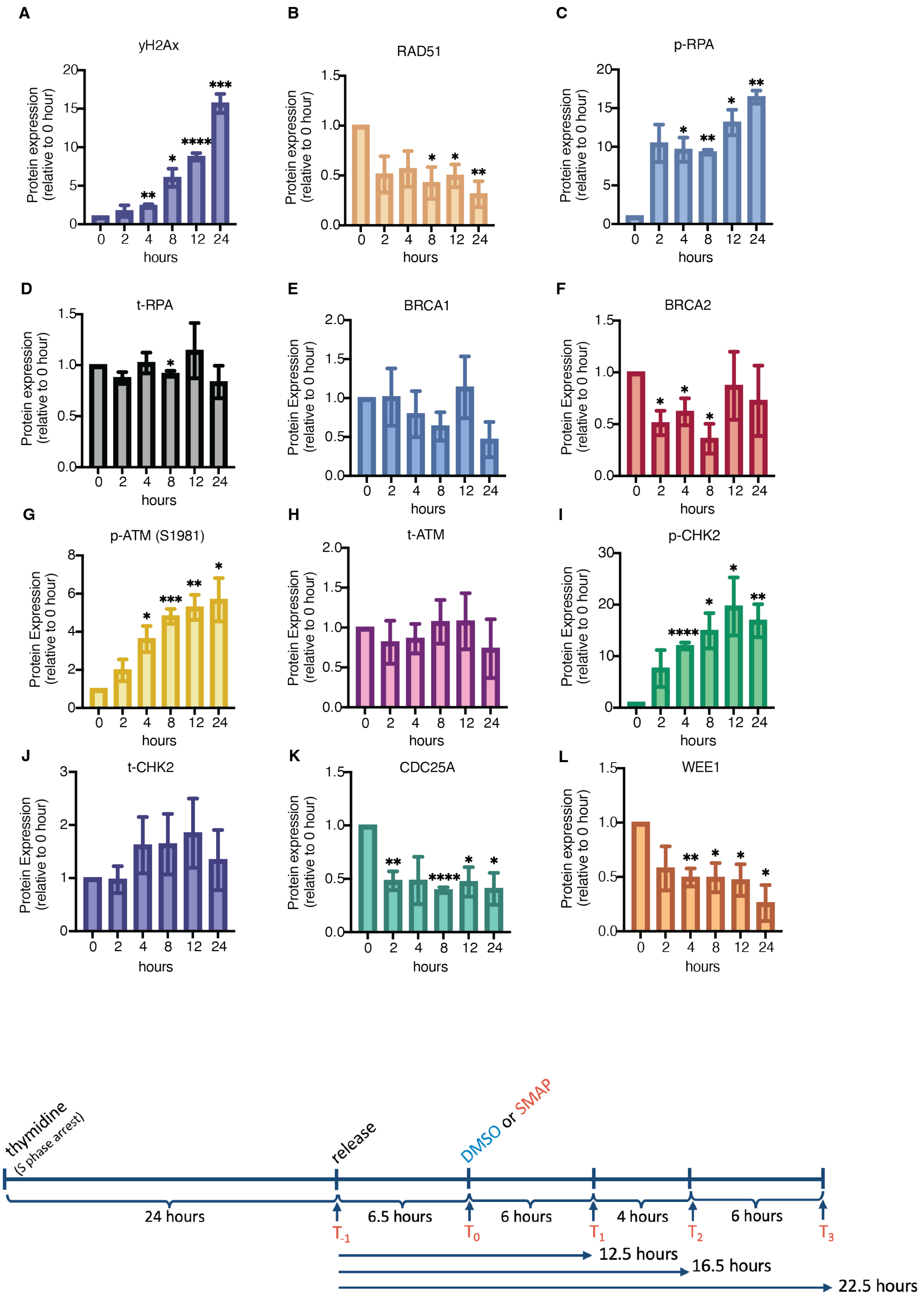

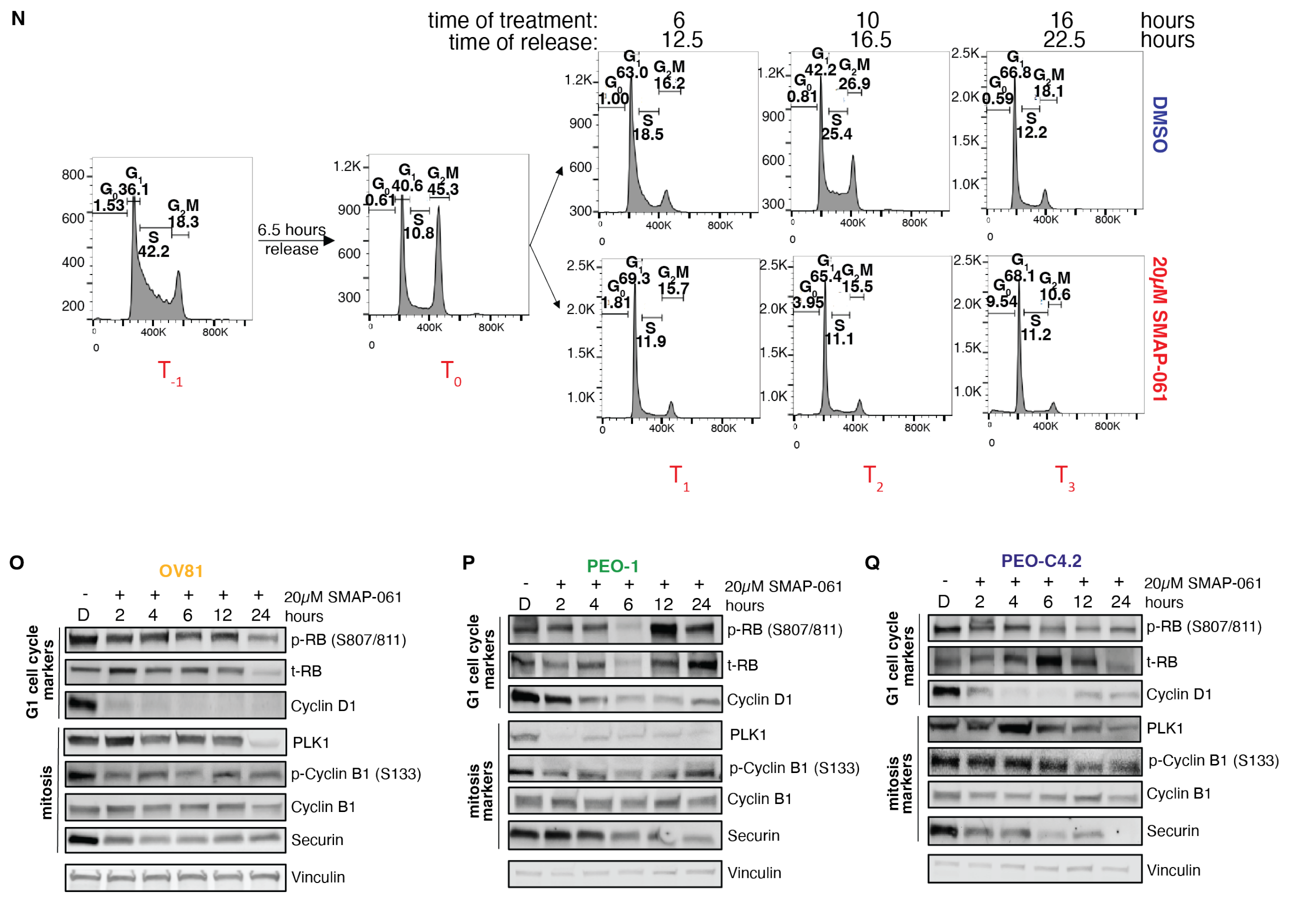


**Supplementary Figure 2 – SMAP-061 consistently decreases expression of DDR and cell cycle proteins in multiple HGSC models, including OV81, PEO-1 and PEO-C4.2, resulting in unrepaired DNA damage and G1 cell cycle arrest.** A) γH2Ax, B) RAD51, C) p-RPA, D) t-RPA, E) BRCA1, F) BRCA2, G) p-ATM, H) t-ATM, I) p-CHK2, J) t-CHK2, K) CDC25A and L) WEE1 protein expression quantification from western blot analysis of OV81 in main Fig. 3B looking at the impact of 20μM SMAP-061 in DDR and HR proteins after 2, 4, 6, 12 and 24 hours of treatment. Data is presented as the mean ± SEM (n=3), (unpaired Student T-tests, comparing each time point relative to zero, *p < 0.05, **p < 0.01, ***p < 0.001, ****p < 0.0001). M) Schematic of the timeline for the treatment and collection of the different time points used for cell cycle flow cytometry analysis. $T_{-1}$ – 24h post 2mM thymidine block; $T_0$ – 6.5 hours post thymidine release; $T_1$ – 6 hours post DMSO or SMAP-061 treatment; $T_2$ – 10 hours post DMSO or SMAP-061 treatment; $T_3$ – 16 hours post DMSO or SMAP-061 treatment. N) Flow cytometry assay of OV81 evaluating cell cycle profiles. Cells were incubated with 2mM thymidine for 24 hours for synchronization in S phase. Thymidine was then released to allow cells to reengage cell cycle progression, by replacing with drug-free media for 6.5 hours. Consecutively, cells were treated with either DMSO (top) or SMAP-061 (bottom) for 6, 10 and 16 hours and cell cycle profiles were analyzed using FlowJo. (Statistics in Fig. 3G). Western blot analysis of O) OV81, P) PEO-1 and Q) PEO-C4.2 cell lines further validating the impact of 20μM SMAP-061 in cell cycle proteins after 2, 4, 6, 12 and 24 hours of treatment.

Supplementary Figure 3

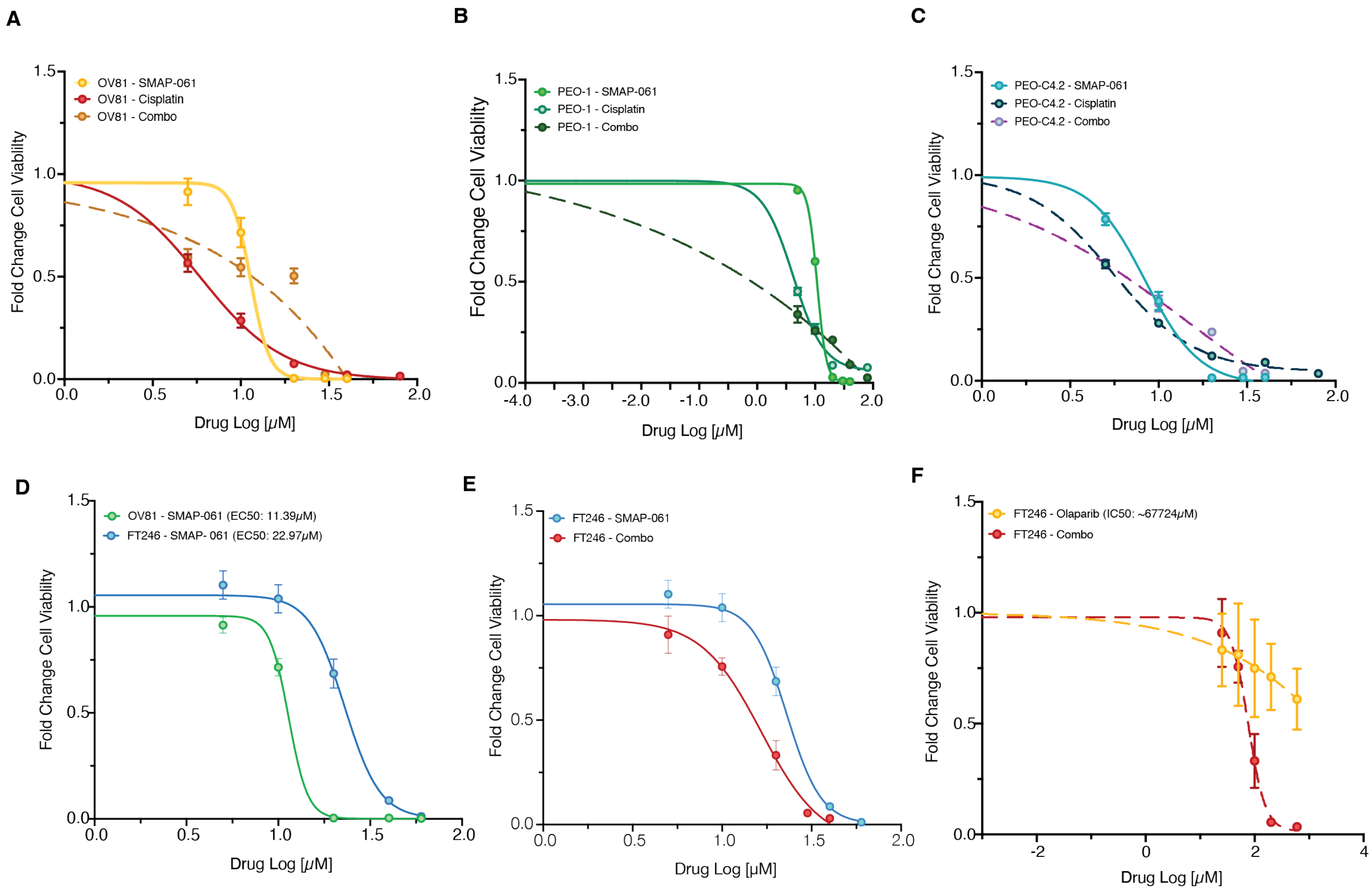

**Supplementary Figure 3 – HGSC and Fallopian Tube non-malignant cells respond differently to SMAP-061 alone and to combination strategies.** A) OV81, B) PEO-1 and C) PEO-C4.2 cells were treated with increasing doses of SMAP-061, cisplatin or combination of both to measure cell viability at 24 hours using MTT. D) Comparison of FT246 and OV81 responses to SMAP-061 utilizing MTT was performed. E) and F) FT246 cells were treated with increasing doses of SMAP-061, Olaparib, or combination of the two and cell viability after 24 hours were measured. Data presented as the mean ± SD (n=3).

Supplementary Figure 4

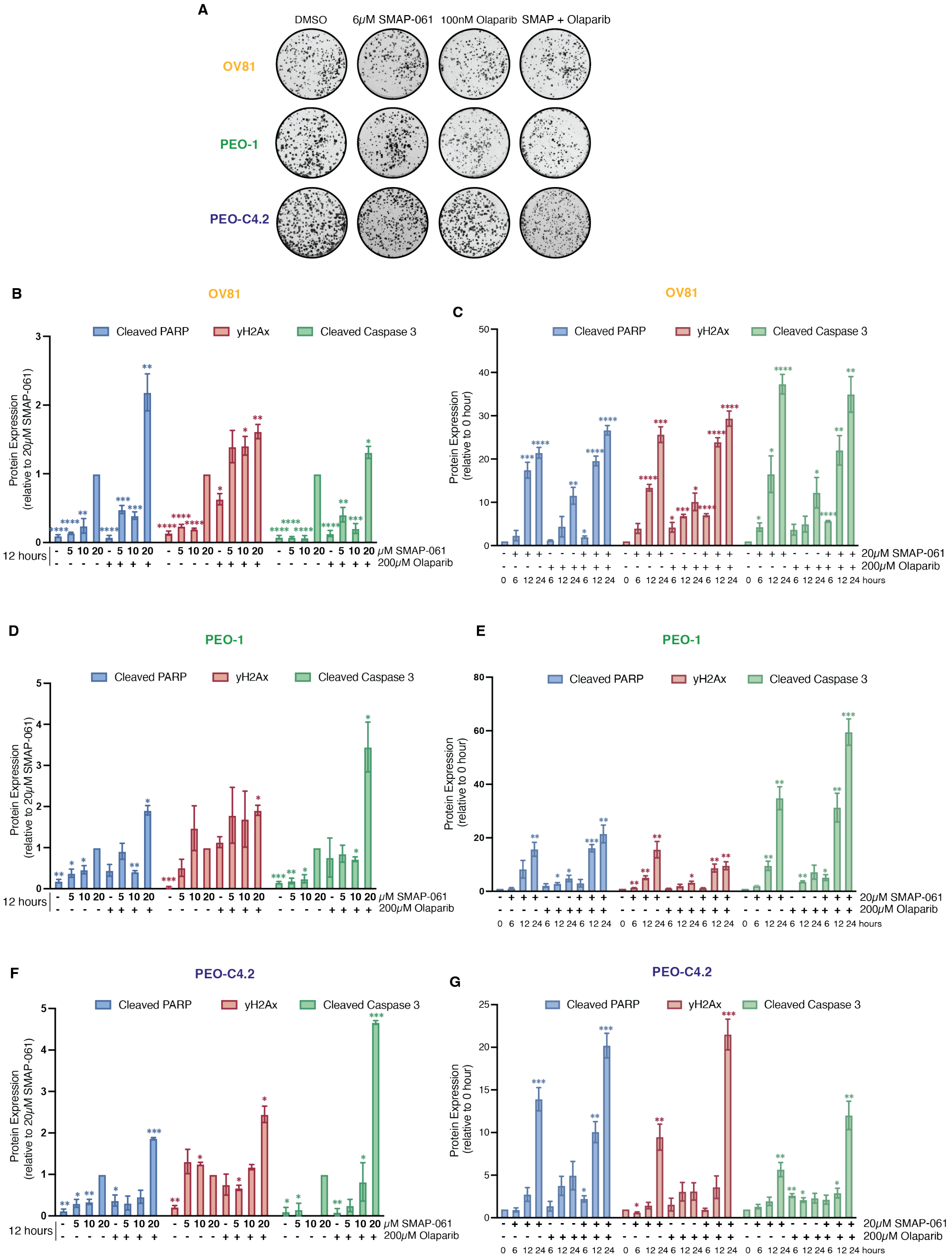

**Supplementary Figure 4 – Combination of SMAP-061 and PARPi synergistically reduces cellular proliferation capacity and induces cell death in a dose and time dependent manner.** A) Clonogenic assay of OV81, PEO-1 and PEO-C4.2 cells treated with DMSO, 6μM of SMAP-061, 100nM of Olaparib or combination of SMAP + Olaparib for the course of two weeks (n=3). B) and C) OV81, D) and E) PEO-1 and G) and H) PEO-C4.2 western blot quantification of cell death markers (Cleaved Caspase 3 and Cleaved PARP) and γH2Ax to evaluate synergy potential of SMAP-061 and Olaparib co-treatment in a dose B), D), F) and time C), E), G) dependent manner. Data is presented as the mean ± SEM (n=3), (unpaired Student T-tests, comparing each condition relative to 20μM SMAP B), D), F) or no treatment at zero hours C), E), G), of each respective cell line, *$p < 0.05$, **$p < 0.01$, ***$p < 0.001$, ****$p < 0.0001$).

Supplementary Figure 5

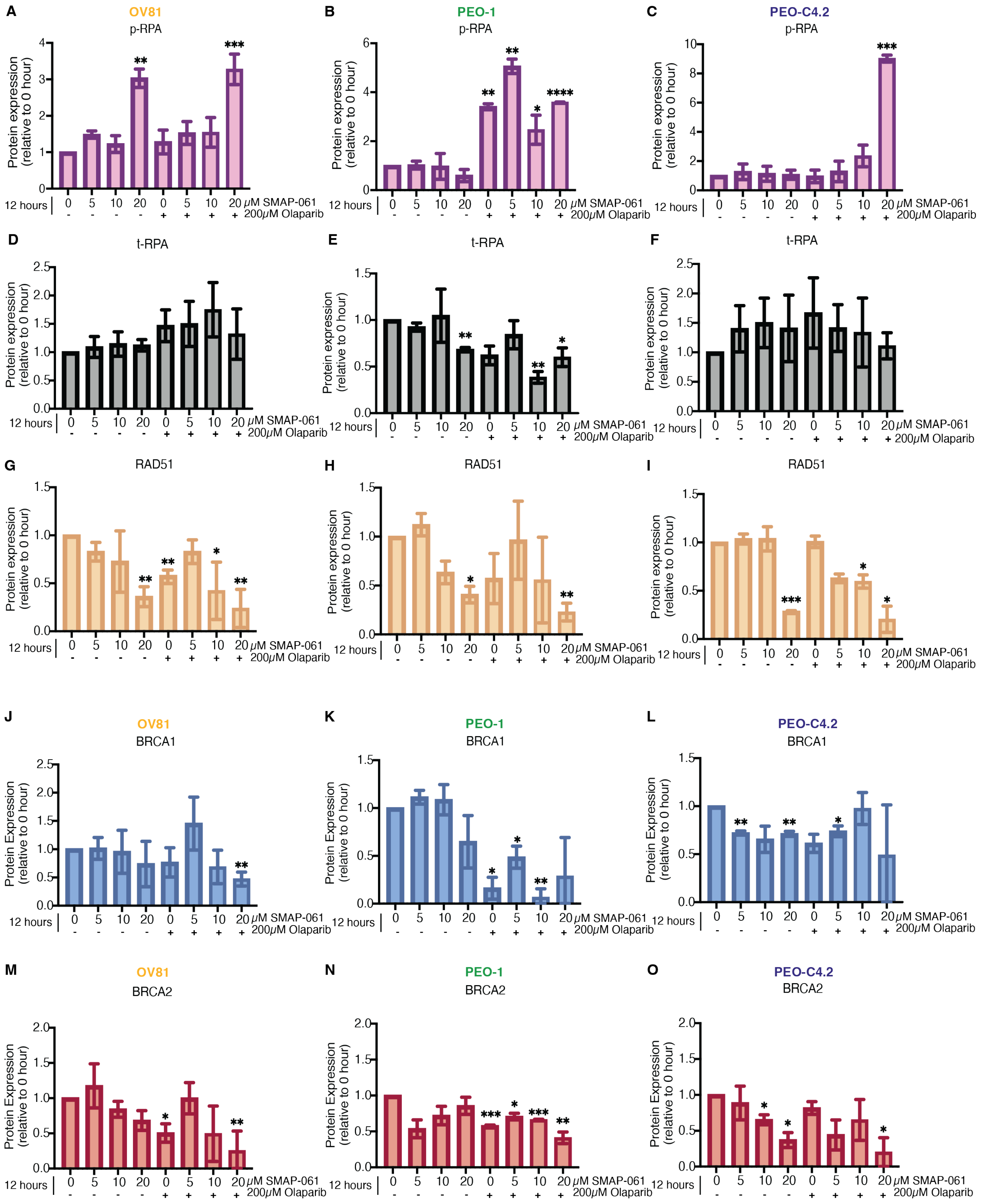

Supplementary Figure 5

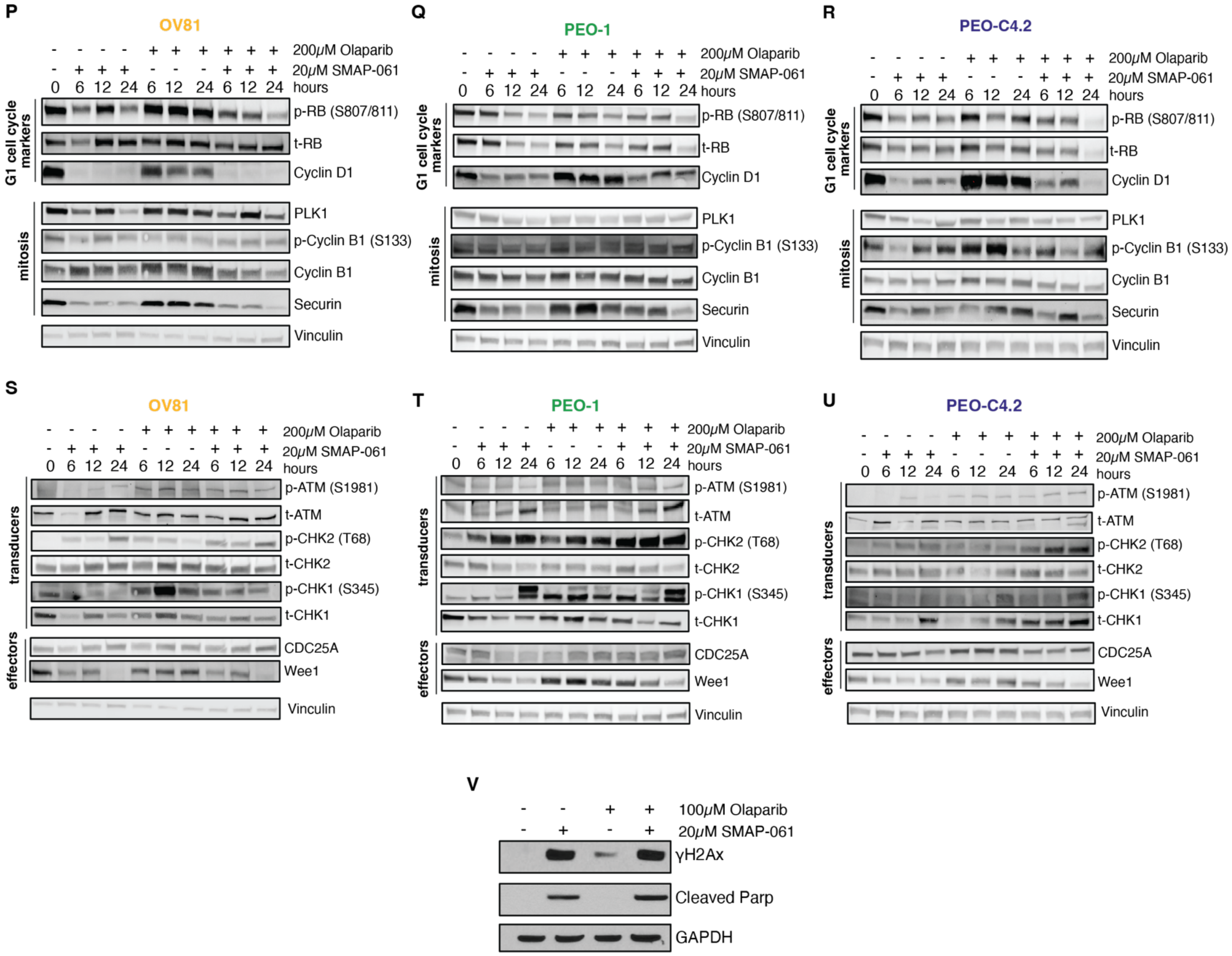


**Supplementary Figure 5 – Co-treatment of SMAP-061 and PARPi synergistically engages the cell cycle and the DDR pathways in a time dependent manner.** A) – O) Quantitation of protein targets expression evaluated on the western blot analysis from Fig. 5A, for OV81, PEO-1 and PEO-C4.2, respectively – A), B), C) p-RPA, D), E), F) t-RPA, G), H), I) RAD51, J), K), L) BRCA1, and M), N), O) BRCA2. Data is presented as the mean ± SEM (n=3), (unpaired Student T-tests, comparing each condition relative to DMSO, *p < 0.05, **p < 0.01, ***p < 0.001, ****p < 0.0001). P)-U) Western blot analysis evaluating cell cycle protein expression regulation upon 20µM SMAP-061, 200µM Olaparib or SMAP + Olaparib treatments for P) OV81, Q) PEO-1 and R) PEO-C4.2 during 6, 12 and 24 hours of exposure. **Transducer** and **effector** proteins (Fig. 3A schematic) are also affected by SMAP-061 treatment and significantly more engaged when in combination with PARPi for S) OV81, T) PEO-1 and U) PEO-C42 cell lines. V) Western blot analysis evaluating Cleaved PARP and γH2Ax expression levels upon 20µM SMAP-061, 100µM Olaparib or SMAP + Olaparib treatments. GAPDH was used as housekeeping.

Supplementary Figure 6

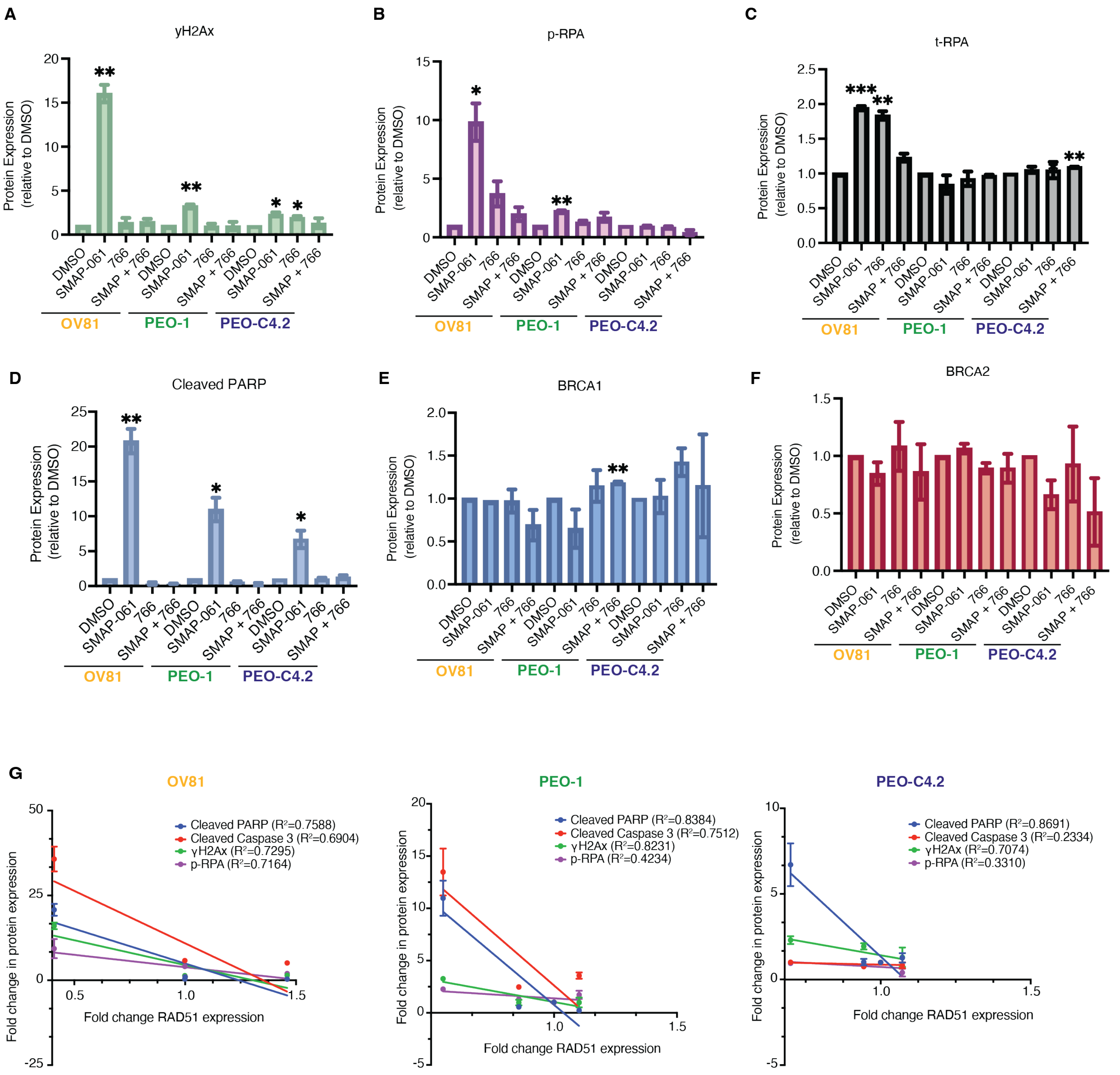

**Supplementary Figure 6 – SMAP-061 specifically targets and downregulates RAD51 through PP2A's activity.** Quantitation of protein targets expression evaluated on the western blot analysis from Fig. 5C, for OV81, PEO-1 and PEO-C4.2, in the presence of DMSO, 20µM SMAP-061, 80µM 766 or SMAP + 766 combination – A) γH2Ax, B) p-RPA, C) t-RPA, D) Cleaved PARP, E) BRCA1, and F) BRCA2. Data is presented as the mean ± SEM (n=3), (unpaired Student T-tests, comparing each condition relative to the DMSO condition of each respective cell line, *p < 0.05, **p < 0.01). G) Correlation analysis graph and $R^2$ values comparing RAD51 protein expression with the expression levels of γH2Ax, p-RPA, cleaved PARP, and cleaved Caspase 3 for OV81, PEO-1 and PEO-C4.2 treated with 20µM SMAP-061. Data presented as the mean ± SEM (n=3).

Supplementary Figure 7

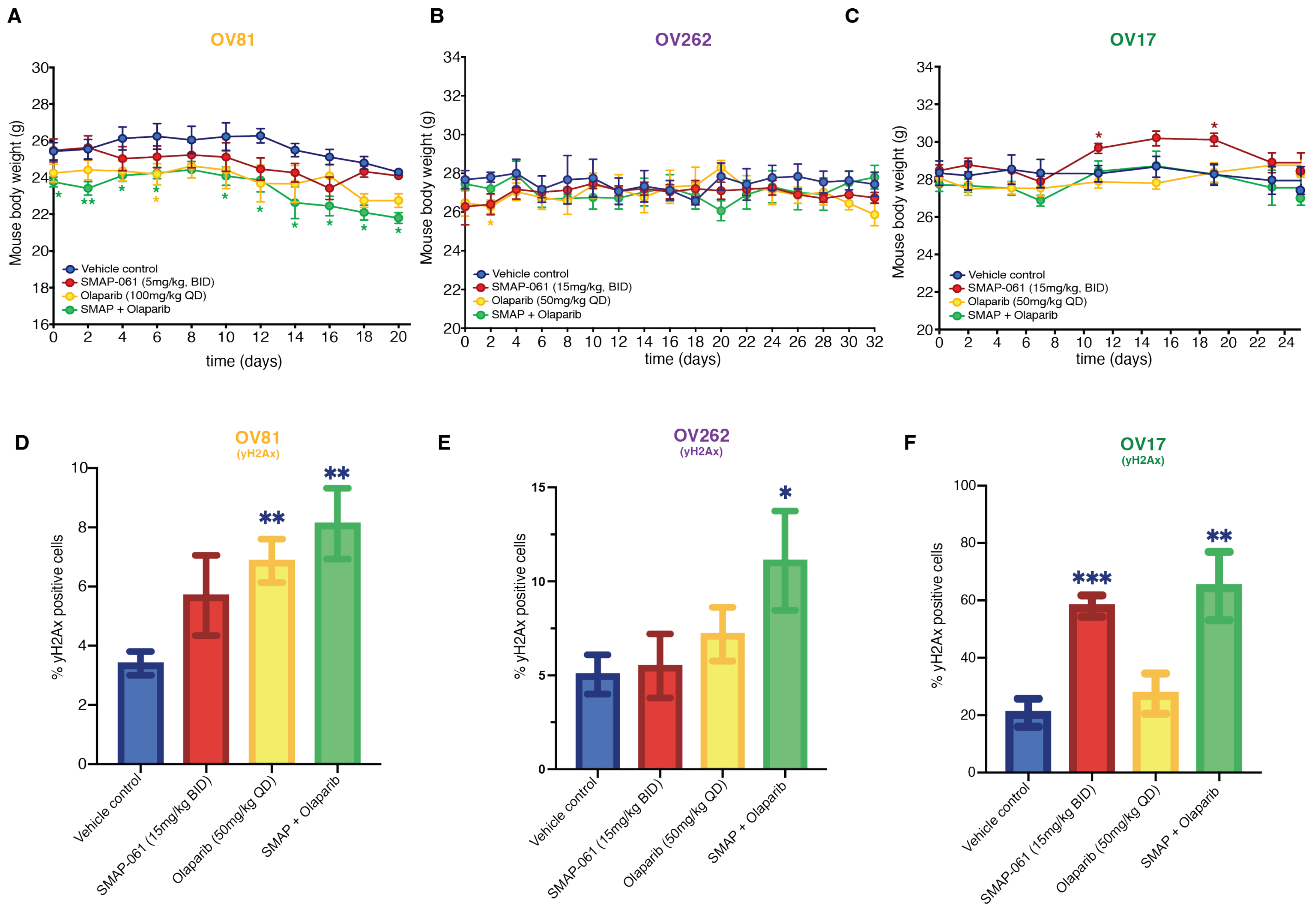

**Supplementary Figure 7 – SMAP-061 show no toxicity effects *in vivo*, having no significant impact on mouse body weight.** A) OV81, B) OV262.2 and C) OV17.1 PDX studies were conducted with tumors implanted in the right flank of NSG mice and allowed to grow between ~80-250mm$^3$ before enrollment in one of 4 treatment groups: Vehicle control, SMAP-061, Olaparib or SMAP + Olaparib combination. Mouse body weights were measured every other day. Data plotted as a function of time and presented as mean ± SEM (Student T-tests, comparing each treatment group relative to vehicle control, *$p < 0.05$, **$p < 0.01$). D) Quantification of *in vivo* $\gamma$H2Ax foci formation for C) OV81, E) OV262 and F) OV17 were calculated from Fig. 6C, 6F and 6I, respectively. Data presented as mean ± SEM (Student T-tests, comparing each treatment group relative to vehicle control, **$p < 0.01$, ***$p < 0.001$).
